# Spatial organization of pulmonary type 2 inflammation by a macrophage-derived cholesterol metabolite

**DOI:** 10.1101/2025.07.29.666625

**Authors:** Yufan Zheng, Hannah E. Dobson, Makheni Jean Pierre, Lilly LeBlanc, Chinaemerem U. Onyishi, Dominic P. Golec, Nathan Carrillo, Anshu Deewan, Claudia A. Rivera, Eduard Ansaldo, Eric V. Dang

**Author notes:** Correspondence (E.V.D).

## Abstract

Effective pulmonary immunity requires the precise spatial organization of immune cells, yet the mechanisms guiding their intratissue positioning during inflammation remain unclear. Here, we identify a cholesterol-derived chemotactic axis that spatially organizes T helper 2 (T_H_2) cells during fungal-induced pulmonary type 2 inflammation. Inflammation-expanded macrophages expressing cholesterol-25-hydroxylase (CH25H) produce 25-hydroxycholesterol, which is converted into the oxysterol 7α,25-dihydroxycholesterol to attract GPR183-expressing T_H_2 cells into infectious lesions. This T_H_2 positioning suppresses interferon-γ responsiveness in inflammatory Ly6C⁺ macrophages, promoting fungal persistence. Disruption of this axis via T_H_2-specific GPR183 deletion restores type 1 macrophage activation and enhances fungal clearance. Our findings reveal a macrophage-driven, metabolite-based mechanism of immunosuppressive cell positioning in inflamed lung tissue.

## Main Text

Effective immune responses in inflamed tissues require spatial coordination of immune cells. This is particularly critical in the lung, where immune responses must be tightly regulated to prevent excessive inflammation and preserve tissue integrity. However, how immune cells are positioned within inflamed lung tissues, and how such positioning contributes to protective or suppressive outcomes, remains poorly understood. Immune cell positioning has been predominantly characterized in lymphoid organs, where G-protein coupled receptor 183 (GPR183; also known as EBI2) serves as a chemotactic receptor that controls immune cell migration to support dendritic cell homeostasis, T cell-dependent antibody responses, and naïve T cell lymph node entry during inflammation (*1–10*). GPR183 ligands are cholesterol metabolites called oxysterols, the most potent of which is 7α,25-dihydroxycholesterol (7α,25-HC). Synthesis of 7α,25-HC from cholesterol requires the stepwise action of cholesterol-25-hydroxylase (CH25H) and oxysterol-7α-hydroxylase (CYP7B1) (*11–14*). At homeostasis, chemotactic 7α,25-HC gradients are generated by non-hematopoietic cells in secondary lymphoid organs, skin, and intestine due to basal constitutive expression of *Ch25h* and *Cyp7b1* by different subsets of stroma (*1, 11, 15–19*).

Macrophages can induce *Ch25h* expression in response to cytokines such as type I/II interferons and interleukin-4 (IL-4)/IL-13 (*14, 20–24*). Interferon-induced *Ch25h* has diverse immunomodulatory functions in macrophages (*25–32*). However, despite being one of the few macrophage IL-4-inducible genes conserved between mice and humans (*20, 33*), the role of *Ch25h* in type 2 immunity is poorly characterized. Most *Ch25h* functions in macrophages have been attributed to the cell-autonomous activity of 25-HC, as macrophages have low expression of CYP7B1 and thus are unlikely to directly synthesize chemotactic sterols (*34*). Despite clear evidence that 25-HC can be secreted (*28, 34, 35*), it remains unknown whether macrophage-derived 25-HC can contribute to 7α,25-HC production in barrier tissues to mediate immune cell migration (*15*).

GPR183 levels in type 2 lymphocytes were reported to be strongly correlated with asthma outcomes following anti-IL-5 antibody treatment (*36*), highlighting the potential importance of GPR183 in regulating pulmonary type 2 immunity. During type 2 responses, T helper 2 (T_H_2) cells and/or group 2 innate lymphoid cells (ILC2s) produce cytokines to defend against parasites and to promote tissue repair (*37, 38*), but can also impair bacterial and fungal clearance (*39–41*). While ILC2s are tissue-resident and can expand locally upon stimulation, T_H_2 cells are initially primed in the draining lymph nodes and then transmigrate from the blood into the lungs during allergic inflammation in response to CCR4 ligands (CCL17, CCL22) (*42, 43*). However, after their tissue entry, how T_H_2 cells achieve proper microanatomical positioning to interact with effector cells within the lung parenchyma remains unknown. Here, we show that GPR183 on T_H_2 cells senses macrophage-derived oxysterols to guide positioning towards infection sites and establish local immunosuppressive niches.

### GPR183 is upregulated on T_H_2 during fungal-induced pulmonary type 2 inflammation

Fungi are potent inducers of type 2 inflammation, with estimates that up to 20% of asthma patients exhibit fungal sensitization (*40*). Notably, type 2 immunity is detrimental during fungal infections, which pose a growing global health concern (*44*). Among these, *Cryptococcus neoformans* (*Cn*) was classified as the highest priority fungal pathogen by the World Health Organization in 2022 (*45*). We infected *Gpr183^Gfp/+^* reporter mice (*46*) intranasally with 5 x 10^4^ *Cn* (KN99α) to induce pulmonary type 2 inflammation and found that ∼60-70 % of lung-tissue resident CD4^+^ T cells expressed GPR183 at 10 days post-infection (dpi), with significantly higher levels than circulating CD4^+^ T cells (Fig. 1, A to C and fig. S1, A to D), implying preferential receptor usage upon tissue entry. Among CD4^+^ T cell subsets, T_H_2 cells had the highest GPR183 expression compared to T_H_1, T_H_17, and regulatory T (T_reg_) cells (Fig. 1, D and E). In vitro polarization of naïve T cells confirmed elevated GPR183 in T_H_2 cells (Fig. 1, F to H). In migration assays, T_H_2 cells exhibited greater sensitivity to the GPR183 ligand 7α,25-HC than T_H_1s, displaying robust chemotaxis toward 0.1 and 1 nM concentrations, whereas T_H_1s only migrated at 10 nM concentrations (Fig. 1, I and J). Importantly, both cell types migrated equally towards 50 nM stromal cell-derived factor 1 (SDF-1) (Fig. 1J). These data link GPR183 upregulation in T_H_2s to increased sensitivity to oxysterol gradients, which may contribute to their microanatomical positioning.

**Fig. 1.**
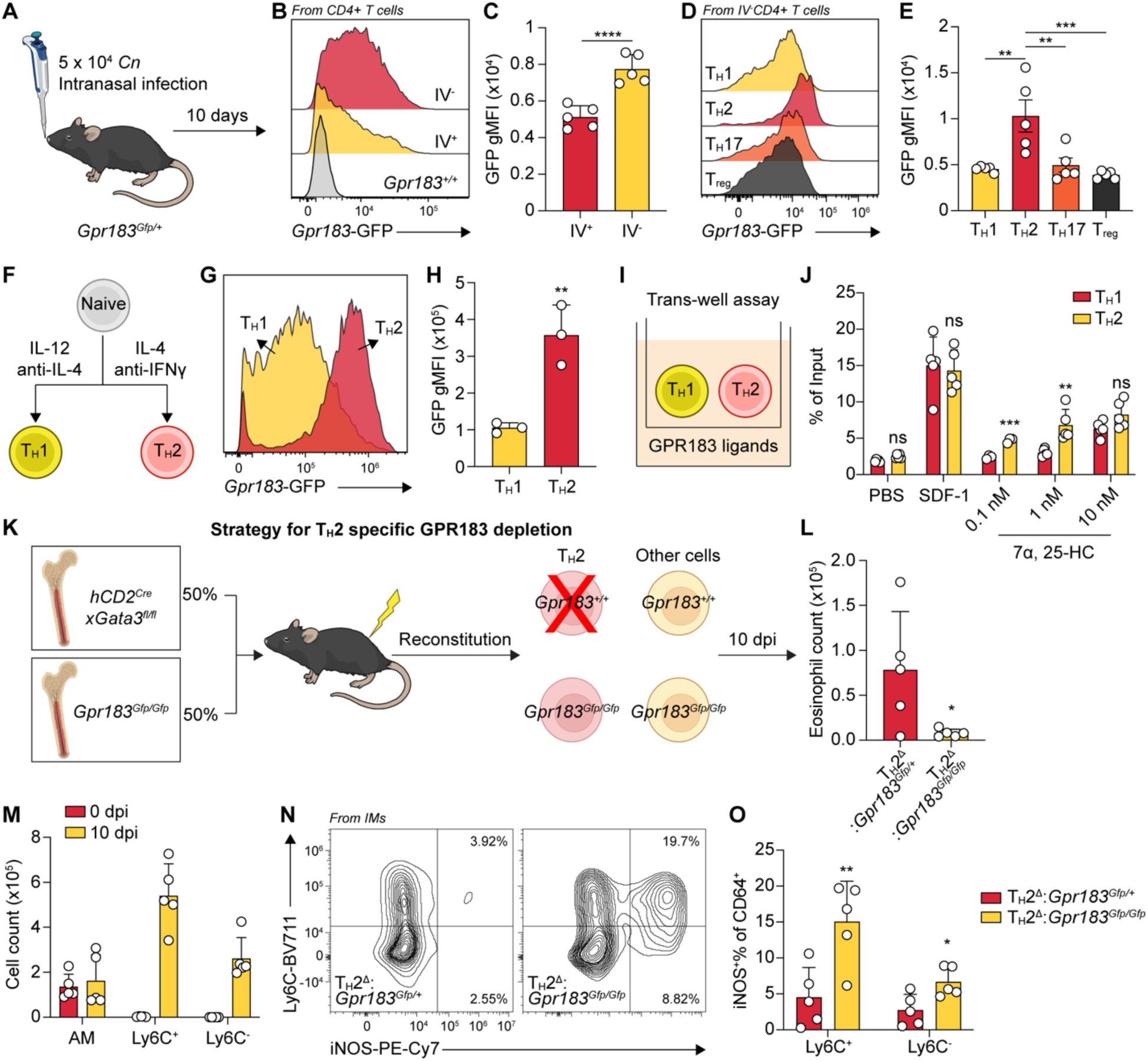
T_H_2 sensing of oxysterols antagonizes iNOS expression in Ly6C^+^ macrophages. (A) Schematic of *Cn* intranasal infection model and intravenous (IV) CD45 labeling. (B) Representative histogram for *Gpr183*-GFP expression in IV^+^ and IV^-^ CD4^+^ T cells from *Cn*-infected mouse lung tissues at 10 dpi. (C) Geometric mean fluorescence intensity (gMFI) of *Gpr183*-GFP in IV^+^ versus IV^-^ CD4^+^ T cells from *Cn*-infected mouse lung tissues at 10 dpi. \*\*\*\**P < 0.0001*, paired two-tailed Student’s t-test. (D) Representative histogram of *Gpr183*-GFP expression across CD4^+^ T cell subsets from *Cn*-infected mouse lung tissues at 10 dpi. (E) *Gpr183*-GFP gMFI in CD4^+^ T cell subsets from *Cn*-infected mouse lung tissues at 10 dpi. \*\**P < 0.01* and \*\*\**P < 0.001* by one-way ANOVA. (F) Schematic of *in vitro* activation protocol for naïve T cells using anti-CD3/anti-CD28 beads with indicated cytokines and blocking antibodies. (G) Representative histogram of *Gpr183*-GFP expression in *in vitro*-generated T_H_1 and T_H_2 cells. (H) Quantification of *Gpr183*-GFP gMFI in *in vitro*-activated T_H_1 and T_H_2 cells. \*\**P < 0.01*, unpaired two-tailed Student’s t-test. (I) Schematic of trans-well migration assay comparing chemotactic responses of *in vitro* activated T_H_1 and T_H_2 cells to GPR183 ligand, 7α, 25-HC. (J) Quantification of T_H_1 and T_H_2 cell migration towards indicated concentrations of 7α, 25-HC and SDF-1. \*\**P < 0.01* and *\*\*\*P < 0.001*; ns, not significant; paired two-tailed Student’s t-test. (K) Schematic of mixed bone marrow chimera strategy for GPR183 specific depletion in T_H_2 cells. (L) Quantification of tissue-resident eosinophils (Live/CD45^+^/IV^-^/CD90.2^-^/ B220^-^/SiglecF^+^/CD11b^+^/SSC-A^hi^) in lungs from *Cn*-infected *Gpr183^TH211^* mice and their controls at 10 dpi. \**P < 0.05*, unpaired two-tailed Student’s t-test. (M) Quantification of lung resident AMs, Ly6C^+^ and Ly6C^-^ IMs at 0 and 10 dpi. (N) Representative flow cytometry plots showing iNOS expression in lung CD64^+^ macrophages from *Cn*-infected *Gpr183^TH211^* mice and their controls at 10 dpi. (O) Quantification of iNOS expression in lung CD64^+^ macrophages from *Cn*-infected *Gpr183^TH211^* mice and their controls at 10 dpi. \**P < 0.05*, \*\**P < 0.01*, unpaired two-tailed Student’s t-test.

### T_H_2 sensing of oxysterols antagonizes iNOS expression in Ly6C^+^ macrophages

To assess the functional role of GPR183 in T_H_2 cells during pulmonary type 2 inflammation, we generated mixed bone marrow chimeras (*hCD2^Cre^xGata3^fl/fl^*:*GPR183^Gfp/Gfp^*), where T_H_2 cells could only derive from *GPR183^Gfp/Gfp^* bone marrow donors because *hCD2^Cre^xGata3^fl/fl^* mice have a selective T_H_2 deficiency (*47*), thus resulting in T_H_2 cell-specific GFP-labeled GPR183 deletion (*Gpr183^TH211^*) (Fig. 1K). At 10 dpi with *Cn*, *Gpr183^TH211^* mice showed reduced eosinophilia (Fig. 1L). Compared to control mice (*hCD2^Cre^xGata3^fl/fl^*:*GPR183^Gfp/+^*), *Gpr183^TH211^* mice showed dramatic increase in inducible nitric synthase (iNOS) expression, particularly in infection-expanded Ly6C^+^ macrophages (Fig. 1, M to O and fig. S1, E and F), suggesting that GPR183 is required on T_H_2 cells to prevent macrophage IFNγ responsiveness. *In vitro*, STAT6 can inhibit IFNγ responsiveness in macrophages by repressing inflammatory gene enhancers (*48, 49*).

However, arginase-1 (ARG1) expression in macrophages was unaltered (fig. S1G), indicating unchanged STAT6 activation. When competed with wildtype macrophages in the same host by mixed bone marrow chimeras, although STAT6-deficient macrophages expressed slightly higher iNOS than wildtype mixing partners, they could not induce iNOS to the same extent as observed in *Gpr183^TH211^* mice (fig. S1, H and I), suggesting only a partial contribution of STAT6 to the suppressed IFNγ responsiveness.

### Lung macrophages establish GPR183 chemotactic gradients via *Ch25h* induction

We utilized a transwell migration assay to assess GPR183 ligand activity in lung tissues, comparing the migration of M12 cells (B lymphoma cell line) transduced with MSCV-*Gpr183*-IRES-*Gfp* versus non-transduced cells towards lung extracts (Fig. 2A). GPR183^+^ cells exhibited significantly greater migration than GPR183^-^ cells towards lung extracts from infected mice at 7 and 10 dpi, but not at 0 or 3 dpi (Fig. 2B), arguing that GPR183 ligand is synthesized *de novo* during infection. While 7α,25-HC is the most potent GPR183 ligand, there is also reported receptor activity for 7α,24-HC (synthesized by CYP46A1 and CYP39A1) and 7α,27-HC (synthesized by CYP27A1 and CYP7B1) (*1, 13, 14, 17*). Of the three enzymes catalyzing the initial cholesterol side chain hydroxylation step, *Ch25h* showed the most robust and sustained induction during infection (Fig. 2C), whereas *Cyp27a1* expression was undetectable and *Cyp46a1* was only transiently induced at 7 dpi (fig. S2A), albeit at ∼1000-fold lower absolute levels than *Ch25h (*fig. S2B). The expression of *Cyp7b1*, responsible for the 7α-hydroxylation step downstream of CH25H and CYP27A1, remained largely unchanged (Fig. 2C). These data suggest that *Ch25h* induction is a key driver of the increased GPR183 ligand activity observed in the lung during infection.

**Fig. 2.**
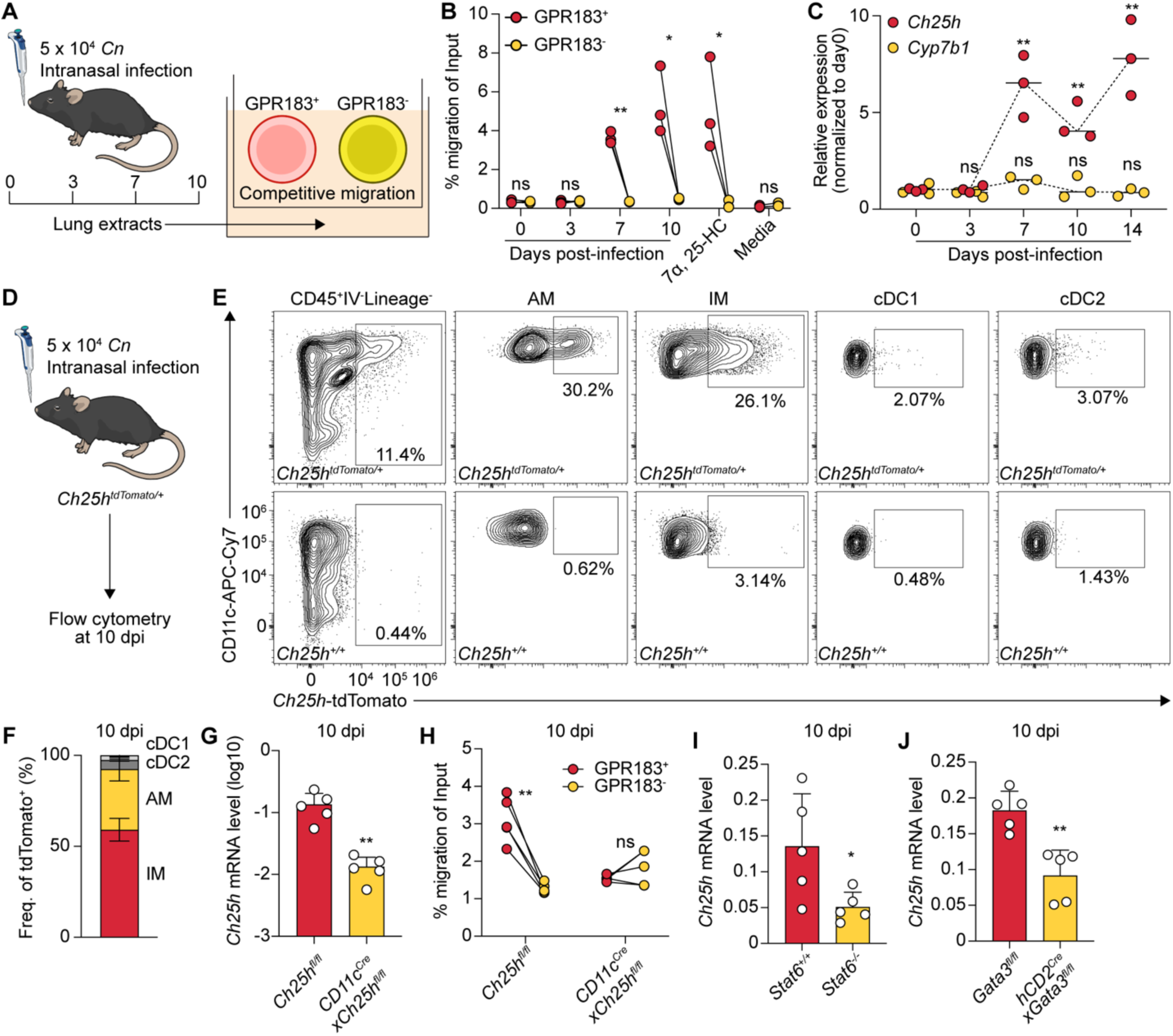
Lung macrophages establish GPR183 chemotactic gradients via *Ch25h* induction. (A) Schematic of trans-well migration assay used to assess GPR183-dependent chemotaxis in lung tissue extracts. (B) Quantification of migration activity of GPR183^+^ versus GPR183^-^ M12 cells in response to lung extracts collected at indicated time points, 1 nM 7α, 25-HC, or migration media alone. \**P < 0.05*, \*\**P < 0.01*; ns, not significant; paired two-tailed Student’s t-test. (C) Relative mRNA levels of *Ch25h* and *Cyp7b1* in lung tissues over time following infection. Expression values were normalized to day 0 to show fold induction. \*\**P < 0.01*; ns, not significant; compared to the expression at day 0 by unpaired two-tailed Student’s t-test. (D) Schematic of *Ch25h*-tdTomato reporter experiments. (E) Representative flow cytometry plots of *Ch25h*-tdTomato expression across indicated immune cell populations from *Cn*-infected mouse lung tissues at 10 dpi. (F) Proportional contribution of each indicated population to total tdTomato^+^ (*Ch25h*-expressing) cells in *Cn*-infected mouse lung tissues at 10 dpi. (G) Relative mRNA levels of *Ch25h* to *Rplp0* in lung tissues from *Cn*-infected *CD11c^Cre^xCh25h^fl/fl^* mice or their controls at 10 dpi. \*\**P < 0.01*, unpaired two-tailed Student’s t-test. (H) Quantification of GPR183^+^ versus GPR183^-^ M12 cell migration towards lung extracts from *CD11c^Cre^xCh25h^fl/fl^* mice and littermate controls at 10 dpi. \*\**P < 0.01*; ns, not significant; paired two-tailed Student’s t-test. (I) Relative mRNA levels of *Ch25h* to *Rplp0* in lung tissues from *Cn*-infected STAT6-deficient mice or their controls at 10 dpi. (J) Relative mRNA levels of *Ch25h* to *Rplp0* in lung tissues from *Cn*-infected T_H_2-deficient mice or their controls at 10 dpi.

Given the known inducible expression of *Ch25h* in macrophages, we hypothesized that lung macrophages might represent the primary source of 25-HC during type 2 inflammation.

Supporting this hypothesis, flow cytometry analysis of *Ch25h^tdTomato/+^* reporter mice (*9*) infected with *Cn* for 10 days revealed that among all hematopoietic cells, *Ch25h* expression was restricted exclusively to CD11c^+^ macrophages including both alveolar macrophages (AMs) and interstitial macrophages (IMs) but not to classical dendritic cells (cDC1s and cDC2s)(Fig. 2, D to F; fig. S2C). To further validate this, we generated *CD11c^Cre^*x*Ch25h^fl/fl^* mice to selectively delete *Ch25h* in CD11c^+^ cells. In these mice, *Ch25h* expression in whole lung was reduced by approximately 10-fold compared to littermate controls at 10 dpi (Fig. 2G), approaching or falling below baseline levels, suggesting that CD11c^+^ macrophages are the major source of *Ch25h* in the lung. Consistently, day 10 post-infection lung extracts from these mice failed to promote GPR183^+^ cell migration, in contrast to extracts from *Ch25h^fl/fl^* mice (Fig. 2H), arguing that macrophages are the principal driver of GPR183 dependent chemotaxis by providing 25-HC that is converted into 7α,25-HC in the lung during infection.

Notably, *Ch25h* induction was reduced in *Stat6^-/-^*and T_H_2-deficient mice, suggesting a partial dependence on T_H_2 cell-mediated type 2 signaling (Fig. 2, I and J). Consistent with this, IL-4 modestly induced *Ch25h* expression in bone marrow-derived macrophages (BMDMs), and this induction was markedly enhanced by *Cn* stimulation (fig. S2D), likely due to upregulation of IL-4 receptor expression (*50*). Consistent with published observations that the SREBP pathway is important for macrophage alternative activation, as 25-HC inhibits SREBP via INSIG1 binding (*51, 52*), *Ch25h*-deficient BMDMs and PEMs showed an elevated ARG1 expression compared to wildtype controls (fig. S2, E to M), supporting a role for *Ch25h* as a negative regulator of macrophage responses to type 2 cytokines. Thus, *Ch25h* is unlikely to promote type 2 immunity by enhancing alternative activation of macrophages.

### Single-cell RNA sequencing reveals distinct transcriptional programs of *Ch25h*-expressing macrophages

Tissue macrophages originate from either embryonic progenitors or adult granulocyte-monocyte progenitor (GMP)-derived monocytes (*53*). At 10 dpi, the majority of Ly6C^+^ macrophages – including ∼80% of Ly6C^+^ IMs and ∼90% of Ly6C^+^ AMs – were *Ms4a3*-tdTomato^+^, indicating a monocyte origin (*53*). In *Ch25h^tdTomato/+^* reporter mice at the same time point, *Ch25h* expression was significantly enriched in Ly6C^-^ IMs and AM compared to the more monocyte-derived Ly6C^+^ population. These findings suggest that *Ch25h* induction is influenced by macrophage ontogeny, with preferential expression in subsets less associated with recent monocyte recruitment.

To better characterize *Ch25h*-expressing macrophages, we performed single-cell RNA sequencing (scRNA-seq) on sorted Ly6C^-^ lung macrophages (MertK^+^CD64^+^Ly6C^-^) from 15 mice across five time points post-*Cn* infection (n=3/time point; fig. S3A). Initial clustering revealed 8 macrophage populations (Mac1-8), including three *Ch25h*-high populations (Mac3, 4, and 6) with *Ch25h* ranking among their top 10 differential expressed genes (fig. S3B). Mac3 and Mac4 shared similar transcriptional profiles and were combined as *Ch25h^+^* macrophages (*Ch25h^+^*Mac). Mac6 expressed high levels of proliferation markers (*Stmn1*, *Top2a*, and *Mki67*) and were annotated as *Ch25h^+^* proliferating macrophages (*Ch25h^+^* pMac) (Fig. 3E and F; fig. S3B). Homeostatic clusters (Mac1, 2, and 5) were categorized into homeostatic macrophages (HoMac) and proliferating homeostatic macrophages (pHoMac) based on proliferative gene signatures (Fig. 3E). Mac7 and Mac8 were classified by their defining markers, *Arg1* and *Mgl2*, respectively (Fig. 3E). *Ch25h^+^* Mac showed elevated expression of *Fabp4* and *Fbp1*, indicating potential metabolic reprogramming, along with *Cdh1*, a marker of alternative activation and epithelioid transformation (*54, 55*) (Fig. 3G). Their abundance peaked at 7 dpi and 10 dpi (Fig. 3H), consistent with peak GPR183-dependent migratory activity.

**Fig. 3.**
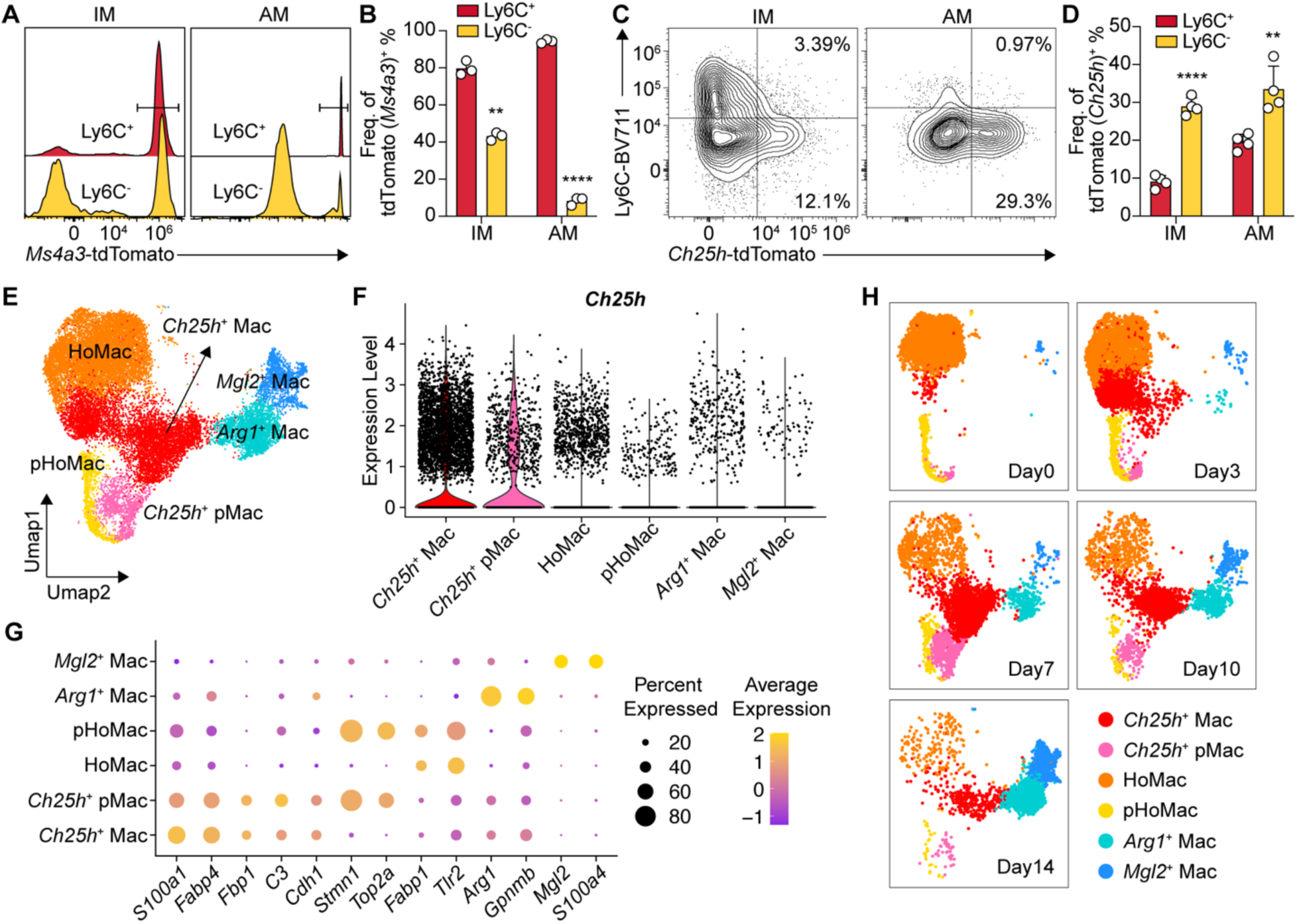
Single-cell RNA sequencing reveals distinct transcriptional programs of *Ch25h*-expressing macrophages (A) Representative histogram of *Ms4a3*-tdTomato expression in Ly6C^+^ versus Ly6C^-^ IMs and AMs from *Cn*-infected mouse lungs at 10 dpi. (B) Ms4a3^+^ frequency of Ly6C^+^ versus Ly6C^-^ IMs and AMs from *Cn*-infected mouse lungs at 10 dpi. \*\*\*\**P < 0.0001* by one-way ANOVA. (C) Representative flow cytometry plot of *Ch25h*-tdTomato expression in Ly6C^+^ versus Ly6C^-^ IMs and AMs from *Cn*-infected mouse lungs at 10 dpi. (D) Frequency of *Ch25h^+^* within Ly6C^+^ versus Ly6C^-^ IMs and AMs from *Cn*-infected mouse lungs at 10 dpi. \*\*\*\**P < 0.0001*; paired two-tailed Student’s t-test. (E) UMAP visualization of annotated macrophage subsets from combined single-cell RNA sequencing replicates (21,069 cells from 15 mice; 3 mice per time point). (F) Violin plot showing *Ch25h* expression across indicated macrophage subsets. (G) Top differentially expressed genes within each macrophage population. (H) UMAP visualization of macrophage subset distribution across individual time points.

Together, these data identify a subset of specialized CD11c^+^ Ly6C^-^ macrophages as the dominant source of inflammation-induced *Ch25h* expression, establishing chemotaxis in lung tissues to promote GPR183^+^ cell migration.

### GPR183 is dispensable for T cell recruitment or differentiation

As GPR183 has been reported to promote eosinophil and macrophage entry to lung tissues in inflammatory settings (*56–60*), we asked whether GPR183 controlled T_H_2 lung entry during fungal infection. At 10 dpi, *Gpr183^TH211^* mice showed no difference in T_H_2 numbers compared to controls (fig. S4A), indicating that GPR183 is dispensable for T_H_2 recruitment. We additionally generated mixed bone marrow chimeras using CD45.1^+^ wildtype and CD45.2^+^ *Gpr183^Gfp/+^* or *Gpr183^Gfp/Gfp^* donors to assess the contribution of GPR183 to tissue entry. At 10 dpi, no differences were observed between the contribution of CD45.2^+^ cells to tissue resident and circulating CD4^+^ T cells, or to lung resident CD4^+^ T cell subsets (fig. S4, B to F), further supporting a recruitment-independent role for GPR183.

As GPR183 has been described to regulate CD4^+^ T cell priming in lymph nodes during ovalbumin immunization (*16*), we also assessed T cell differentiation in GPR183-knockout mixed bone marrow chimeras in both lung tissues and lung-draining mediastinal lymph nodes. At 10 dpi, GATA3, T-bet, and RORγT expression, as well as cytokine production upon restimulation, were equivalent in lung-resident CD4^+^ T cells (fig. S4, G to O) and no differences were found in GATA3 or T-bet expression in CD4^+^ T cells in lymph nodes (fig. S4, P and Q), indicating that GPR183 does not affect CD4^+^ T cell differentiation in the setting of fungal infection.

To test whether any effects of GPR183 on T_H_2 recruitment or differentiation were being masked in the polyclonal response, we generated *Cn* chitin deacetylase 2(CDA2)-specific TCR transgenic (CnT II) mice (*61, 62*) that were either *Gpr183^Gfp/+^* (CD45.1) or *GPR183^Gfp/Gfp^*(CD45.1/2) and performed competitive adoptive CD4^+^ T cell transfers into *Ch25h^fl/fl^* or *CD11c^Cre^*x*Ch25h^fl/fl^* mice (CD45.2) (fig. S5, A and B). At 10 dpi, the ratio of transferred *Gpr183^Gfp/+^* to *GPR183^Gfp/Gfp^*T cells in the lung was identical between *Ch25h^fl/fl^* and *CD11c^Cre^*x*Ch25h^fl/fl^* mice (fig. S5C), and the GATA3^hi^ % was equivalent between *Gpr183^Gfp/+^*and *GPR183^Gfp/Gfp^* transferred FOXP3^-^CD4^+^ T cells (fig. S5D) in both recipients, further supporting a dispensable role for GPR183 in fungal antigen-specific CD4^+^ T cell recruitment and T_H_2 differentiation.

### GPR183 orients T cells towards fungal granulomatous lesions

Given that GPR183 deficiency did not affect T cell recruitment or differentiation, we hypothesized that it regulates intrapulmonary positioning. As wildtype KN99α *Cn* infection typically causes diffuse lung infection that makes it difficult to resolve positioning differences, we utilized a granuloma-forming strain (*41, 63*), *Cn gcs1Δ* (5 x 10^4^ intranasal infection), to spatially observe immune cell positioning within organized infectious lesions. We first confirmed no T cell recruitment or differentiation effects by GPR183 in this model at 21 dpi (fig. S6, A to C), when the mice had the highest T_H_2 cell numbers within the infection time course (*41*).

To test whether GPR183 regulates parenchymal immune cell positioning, we overexpressed GPR183 in hematopoietic cells by transducing a MSCV-*GPR183*-*Gfp* retroviral vector into bone marrow, which was then transplanted into irradiated recipients. At 35 dpi, the peak of pulmonary fungal load when organized structures start being observed, by thick tissue confocal imaging, we found that GPR183-overexpressing cells were all localized with the granulomatous area at 35 dpi (defined by CD11c staining and *Cn* mCherry signals), whereas a large portion of control GFP-transduced immune cells were still around vasculature (defined by autofluorescence) (Fig. 4A). Quantitative spatial analysis showed a bimodal distribution of control GFP^+^ cells: one peak near the vasculature and another one at the infection site represented by high density of mCherry signals, while GPR183-overexpressing cells only displayed a single peak colocalized with mCherry peak-infection sites (fig. S6D), demonstrating that GPR183 overexpression could force immune cell migration into granulomatous lesions.

**Fig. 4.**
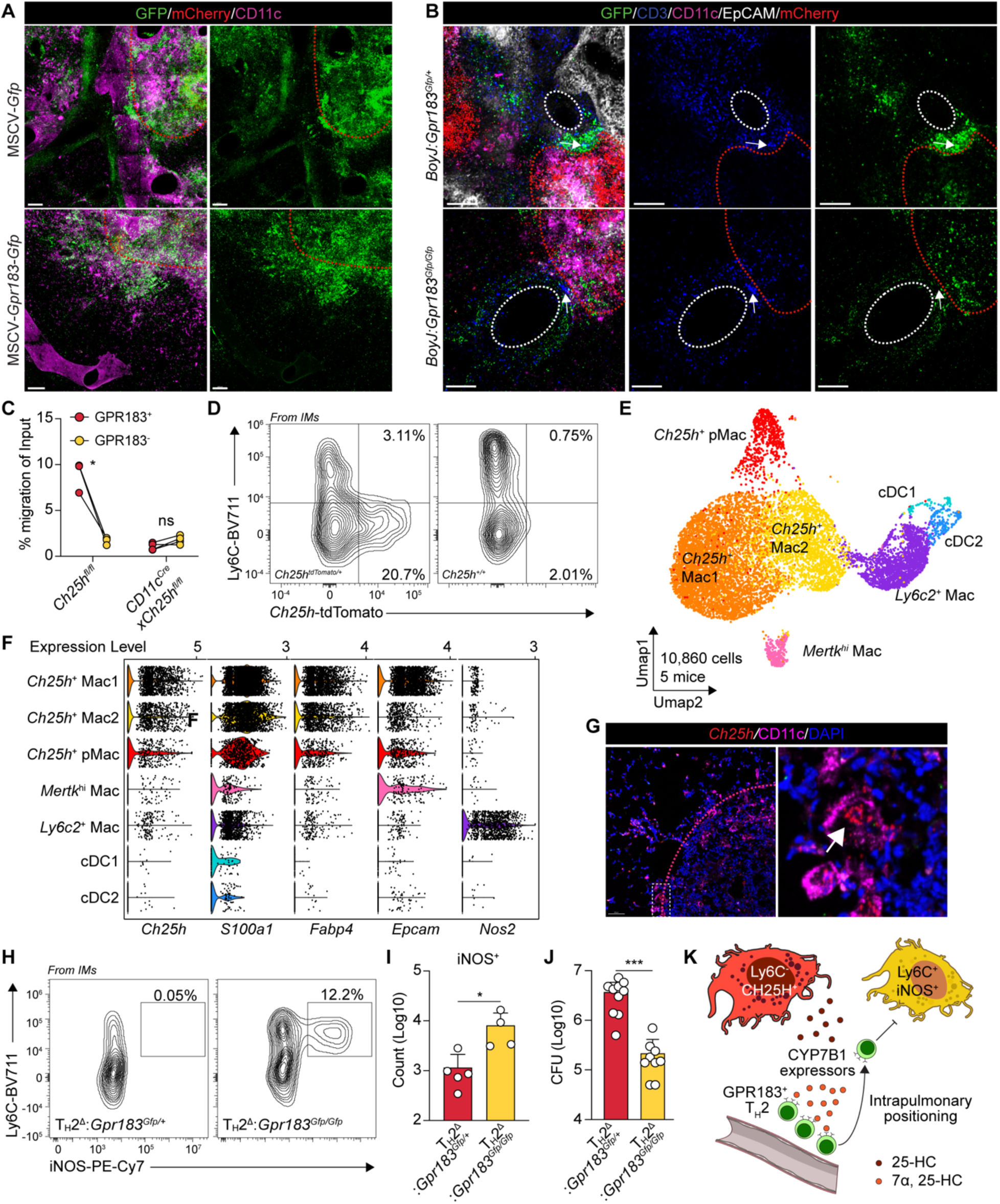
GPR183 guides intrapulmonary positioning of T_H_2 cells to antagonize fungal clearance within granulomatous lesions. (A) Representative full-projection confocal images of 500 μm thick lung sections from bone marrow chimeras overexpressing MSCV-*Gfp* or MSCV-*Gpr183*-*Gfp* from *Cn gcs1Δ*-infected at 35 dpi. (B) Representative full-projection confocal images of 500 μm thick lung sections from *Cn gcs1Δ*-infected *Gpr183*-knockout mixed bone marrow chimeras or their controls at 35 dpi. (C) Quantification of migration activity of GPR183^+^ versus GPR183^-^ M12 cells in response to lung extracts collected from *CD11c^Cre^xCh25h^fl/fl^* mice or their controls at 35 dpi. \**P < 0.05*; ns, not significant; paired two-tailed Student’s t-test. (D) Representative flow cytometry plots of *Ch25h*-tdTomato expression in lung CD64^+^ macrophages at 35 dpi. (E) UMAP visualization of annotated myeloid cell subsets from merged scRNA-seq replicates (10,860 cells from 5 mice at 35 dpi). (F) Expression patterns of selected genes across indicated myeloid subsets. (G) Representative imaging of *Ch25h* transcripts detected by RNAscope, co-stained with CD11c by immunofluorescence on thin sections of *Cn gcs1Δ*-infected mouse lung tissues at 35 dpi. (H) Representative flow cytometry plots showing iNOS expression in CD64^+^ lung macrophages from *Cn gcs1Δ*-infected *Gpr183^TH211^* mice and their controls at 21 dpi. (I) Quantification of iNOS^+^ lung macrophages in *Cn gcs1Δ*-infected *Gpr183^TH211^* mice and their controls at 21 dpi.(J) Pulmonary fungal burden in *Cn gcs1Δ*-infected *Gpr183^TH211^* mice and their controls at 21 dpi. CFU, colony forming unit. (K) Schematic for the proposed model. During inflammation, Ly6C^+^ macrophages induce *Ch25h* expression, leading to the production of oxysterols that establish a GPR183-mediated chemotaxis. This guides T_H_2 intrapulmonary positioning, which in turn antagonize local IFN-γ responses.

Next, we performed similar thick section imaging on lung tissues derived from mixed bone marrow chimeras (*BoyJ*:*Gpr183^Gfp/+^* and *BoyJ*:*Gpr183^Gfp/Gfp^*) with CD3 co-staining to directly ask whether oxysterols could direct T cell positioning through GPR183. We found that generally, there was one group of CD3^+^ T cells around vasculature and another group aggregating towards granulomatous lesions at 35 dpi (Fig. 4B). In *BoyJ*:*Gpr183^Gfp/+^* chimeras, GFP^+^ GPR183-expressing T cells were oriented towards the granulomatous lesions, which was not observed in *BoyJ*:*Gpr183^Gfp/Gfp^* chimeras, where the GFP^+^ GPR183-deficient cells were distributed uniformly around the vasculature (fig. S6E). These data, together with the GPR183-overexpression data, demonstrate that T cells use GPR183 to orient towards granulomatous lesions within the inflamed lung after tissue entry.

### Granuloma-associated macrophages express *Ch25h* during persistent fungal infection

CH25H expression is found within human and mouse mycobacterial granulomas (*64*). As we observed macrophage-derived oxysterols contributing to GPR183 ligand generation during acute infection, we asked whether this also occurs within fungal granulomatous infection. Migration assays showed an increase in GPR183 ligand activity from lung extracts from *Cn gcs1Δ* infected mice across a time course (fig. S7A). *Ch25h* specific deletion in CD11c^+^ cells also abolished the GPR183-dependent migration activity at 35 dpi (Fig. 4C), suggesting a similar group of macrophages expressed *Ch25h* during fungal granulomatous infection that was found during acute infection. Consistently, we found the same enrichment of *Ch25h* signal in CD11c^+^ Ly6C^-^ macrophages at 35 dpi using *Ch25h^tdTomato/+^* reporter mice (Fig. 4D and fig. S7B).

We next performed scRNA-seq on broad myeloid cells (CD64^+^MHCII^+^CD11c^+^) to better characterize *Ch25h^+^* macrophages during fungal granulomatous infection at 35 dpi. Initial clustering identified seven populations (C1 to C7) (fig. S7C). Two clusters (C6 and C7) expressed high *Zbtb46* and were annotated as cDC1 (*Itgae^+^*) and cDC2 (*Mgl2*^+^). The remaining clusters (C1 to C5) expressed *Mertk*, consistent with macrophage identity. C4 with the highest *Mertk* expression, was named *Mertk*^hi^ Mac. Although C5 had lower *Mertk*, it expressed *Mafb*, a transcription factor favoring macrophage over DC differentiation, and was named *Ly6c2*^+^ Mac (fig. S7D).

*Ch25h* expression was enriched in macrophage clusters-C1, C2, and C3 (fig. S7D). *Ly6c2*^+^ Mac did not express *Ch25h*, consistent with the reporter data. C1, expressing proliferative markers (fig. S7D), was named as *Ch25h*^+^ pMac, while C2 and C3 were subsequently named as *Ch25h*^+^ Mac1 and *Ch25h*^+^ Mac2 (Fig. 4E). Notably, *S100a1* and *Fabp4* were consistently co-expressed with *Ch25h* within these clusters. While *Cdh1* expression was low compared to acute infection, *Ch25h*^+^ Mac1 and pMac expressed *Epcam*, an epithelial marker associated with macrophage epithelioid transformation (Fig. 4F). To localize *Ch25h* spatially, we performed *Ch25h*

RNAscope *in situ* hybridization with CD11c co-staining and found that *Ch25h* signals were enriched in large-body CD11c^+^ cells at the granuloma border (Fig. 4G).

### T_H_2 sensing of oxysterols antagonizes fungal clearance within granulomatous lesions

Consistent with wildtype KN99α *Cn* infection, during KN99α *Cn gcs1Δ* infection, *Gpr183^TH211^* mice showed the same increase in iNOS expression in Ly6C^+^ macrophages while ARG1 still remained unchanged (Fig. 4, H and I; fig. S7E) at 21 dpi. Notably, scRNA-seq showed an enrichment of *Nos2* in *Ly6c2*^+^*Mafb^+^*Mac in wildtype mice (Fig. 4F), suggesting Ly6C^+^ macrophages were the main IFNγ signaling responders. A protective role for IFNγ has been described in chronic *Cn* infection (*41, 65–70*). Therefore, we asked whether the increase of IFNγ responsiveness corresponded to a lower pulmonary fungal load. Indeed, *Gpr183^TH211^* mice showed significantly lower pulmonary fungal burden than controls at both 21 and 35 dpi (Fig. 4J and fig. S7F).

In summary, our data suggested a model where Ly6C^-^ *Ch25h*-expressing macrophages produce 25-HC to establish chemotactic gradients that guide T_H_2 positioning towards infectious lesions, thus suppressing iNOS production in Ly6C^+^ macrophages and promoting persistent fungal infection (Fig. 4K).

## Discussion

This study identifies a critical mechanism for T_H_2 positioning within the inflamed lung. Although chemokines are known to regulate cell trafficking from the blood into the lung parenchyma, identification of chemotactic receptors that uncouple recruitment from post-entry positioning has been challenging. Here, we provide direct evidence that spatial localization, independent of recruitment, is a distinct and essential layer of immunoregulation during pulmonary type 2 inflammation. In inflamed lung tissues, GPR183, a well-established chemotactic receptor in lymphoid organs, guides recruited T cells towards infection sites, thereby shaping local immune responses.

Langerhans cells have been shown to partially contribute to GPR183-dependent naïve lymphocyte recruitment into inflamed lymph nodes (*7*), suggesting that inflammation-dependent *Ch25h* expression in myeloid cells can contribute to tissue entry. Here, we identify CD11c^+^Ly6C^-^ lung macrophages that emerge during pulmonary fungal infection as major 25-HC producers establishing GPR183-dependent chemotaxis to guide T_H_2 positioning. Notably, *Cyp7b1* does not significantly increase upon infection, suggesting that *Ch25h* induction in macrophages is rate-limiting for production of chemotactic oxysterols in the lung. Distinct from what has been shown with other cell types (*5–9, 71*), we did not observe a requirement for GPR183 in T cell tissue entry, indicating either redundancy or lack of function in this process. We found that STAT6-or T_H_2-deficient mice have reduced *Ch25h* expression at the whole lung level, suggesting a contribution from type 2 cytokines to *Ch25h* induction, potentially in cooperation with interferon signaling. *Ch25h*^+^ macrophages display elevated lipid metabolism signatures in scRNA-seq data, which may contribute to *Ch25h* upregulation or 25-HC synthesis. In line with previous findings (*72*), homeostatic AMs express baseline levels of *Ch25h*, but we observed little GPR183-dependent migration activity in lung extracts from uninfected mice. This implies that AM-derived 25-HC may either have alternative functions or spatial distributions insufficient to establish chemotactic gradients in cooperation with functional *Cyp7b1*^+^ cells, which are likely adventitial fibroblasts based on expression data from different lung cell types (*73*).

Our findings also offer mechanistic insights into granuloma organization. Beyond cellular recruitment, there must exist spatial cues within tissues that guide immune cell positioning towards granulomatous lesions. We identify a non-redundant role for GPR183-mediated chemotaxis in organizing T_H_2 cells within these microanatomical structures that emerge during chronic fungal infection. Specifically, we show that *Ch25h*^+^ macrophages, expressing markers of epithelioid transformation, guide T_H_2 localization through GPR183. This regulation might be conserved in granulomatous structures across different infectious and non-infectious settings.

Supporting this idea, CH25H expression has been reported around human tuberculosis granulomas (*64*). T lymphocyte cuffs are a common feature of granulomas (*74, 75*), resembling tertiary lymphoid structures that can support both pathogen clearance and persistence. Our study suggests that granuloma-associated macrophages actively shape this niche by directing T_H_2 positioning, thereby promoting local immunosuppression and pathogen persistence.

Additionally, our study sheds new light on the mechanisms underlying type 1 and type 2 immune competition during mixed inflammation. A tempting hypothesis is that T_H_2 cells outcompete T_H_1 cells for oxysterol gradients to co-localize with Ly6C^+^ macrophages, thereby suppressing local type 1 response. Consistent with this idea, GPR183-deficient T_H_2 cells failed to restrain IFNγ responsiveness in macrophages, despite normal tissue T_H_2 numbers, highlighting the importance of spatial proximity for this regulatory interaction. Importantly, this effect is unlikely to be explained solely by loss of STAT6-dependent competition with STAT1. These findings point to a proximal, spatially constrained crosstalk, where Ly6C^-^ macrophages guide T_H_2 cells to Ly6C^+^ macrophages to potentially ‘shield’ them from T_H_1s, thus preventing iNOS induction.

The potential importance of GPR183 has recently been suggested by a clinical study in asthma patients, which found that the therapeutic efficacy of anti-IL-5 treatment correlated with reduced GPR183 expression on type 2 lymphocytes (*36*). Although we did not observe a lung entry effect for GPR183 on T_H_2 cells, our findings demonstrate that GPR183-mediated chemotaxis is essential for their intra-parenchymal migration to inflammation sites, thereby shaping the local immune responses. From a translational perspective, manipulating this spatial axis could offer therapeutic strategies for diseases involving type 1/2 immune imbalances.

## Acknowledgement

We thank Drs. Jason Cyster and Andrea Reboldi for gifting *Gpr183^Gfp/Gfp^*and *Ch25h^tdTomato/+^* mice and *Gpr183*-overexpressed M12 cell line. We thank the NIAID animal facility staff, as well as O. Schwartz and S. Ganesan (NIAID Biological Imaging Facility). We thank Katrin Mayer-Barber, Niki Moutsopoulos, Hao Jin, Dan Barber, and Mihalis Lionakis for helpful discussions.

## Authors contribution

Y.Z. and E.V.D. designed the study and experiments and wrote the original manuscript.

Y.Z. performed the experiments, analyzed the data, and created the figures. Y.Z., H.E.D, M.J.P., C.U.O., and N.C. performed flow cytometry experiments. L.L. performed the migration assays and macrophage *in vitro* cultures. D.G. performed the T cell differentiation *in vitro*. Y.Z. and A.D. performed single cell RNA-seq analysis. Y.Z., C. A.

R. and E.A performed the cell sorting.

## Data and materials availability

All source data are available upon request.

## Supplementary Materials

## Materials and Methods

### Animal

All mouse experiments were approved by the National Institutes of Allergic and Infectious Diseases Animal Care and Use Committee (NIAID-ACUC) and were performed in accordance with NIAID-ACUC guidelines and under approved protocols (LHIM-4E). Female and male mus musculus were used in this study. In bone marrow chimera experiments, animals were irradiated at 6-to 8-week-old and analyzed 6-8 weeks following irradiation. Most other experiments were performed on adult animals between 8 and 20 weeks of age. *Gpr183^Gfp/Gfp^* (*1*) and *Ch25h^tdTomato/+^* (*2*) mice were gifted by Dr. Jason Cyster and Dr. Andrea Reboldi. *hCD2^Cre^xGata3^fl/fl^* mice (line 20939) and *BoyJ* mice (8478) were ordered from NIAID-Taconic Exchange program. *Ch25h^fl/fl^*(JAX:037647) and *CD11c^Cre^* (JAX:008068) mice were purchased from Jackson Laboratory. All mice were housed in a specific pathogen–free environment.

### Bone marrow chimera

Mice were irradiated at 700 rads in two doses spaced 3hrs apart by 14DNR at NIAID. Bone marrow was prepared and injected the same day at least one hour after the final irradiation dose. Mice were monitored during 6-week reconstitution. Bleeding was performed to determine reconstitution efficiency.

### Flow cytometry

All mice for flow cytometry analysis in this study received retroorbital injection of 2 μg CD45 antibody 3 to 5 minutes before euthanasia to label intravascular cells. For lung samples, tissues were collected directly into 5 ml digestion buffer (HBSS with 1 mg/ml collagenase II (Gobio #17-101-015), 1 mg/ml Dispase (Gibco #17-105-041), and 10 μg/ml DNase I (Sigma-Aldrich #11284932001) and then digested using a GentleMACS (Miltenyi Biotec). Digestion was stopped by adding 5 ml flow buffer (PBS with 2% FBS and 2 mM EDTA) on ice. Samples were immediately filtered using 70 μm strainers and centrifuged to remove digestion buffer. Red blood cells were removed by ACK lysis buffer (BioLegend #420302) for 5 min on ice and cells were then spun, washed, and aliquoted in 96-well round bottom plates for staining. Flow antibodies used in this study are listed in Table S1. Live/dead staining was performed for 10 min in PBS containing Fc Block before protein staining. Surface proteins were stained for 20 min on ice and then cells were acquired using a Cytek Aurora spectral flow cytometer. Intracellular and intranuclear proteins were stained with Transcription Factor Staining Buffer set (eBioscience #00-5523-00). Briefly, cells were fixed and permeabilized on ice for 1 hour and stained in permeabilization buffer containing antibodies for 1 hour at room temperature. Cells were washed and resuspended in flow buffer for acquisition. T cell restimulation was performed for cytokine analysis. Collected cells were cultured in RPMI 1640 culture medium with PMA (50 ng/ml) and ionomycin (500 ng/ml) for 4 hours and then processed for staining. Data were analyzed using FlowJo 10.10.0.

### T cell culture in vitro and competitive migration assay

Spleens and Lymph nodes were harvested from mice and mashed through a 100 µm strainer in lymphocyte media (RPMI 1640 containing 10% FBS, 10mM HEPES, 2mM L-Glutamine, 55 µM 2-Mercaptoethanol, and antibiotics). CD4 T cells were isolated using EasySep™ Mouse CD4+ T Cell Isolation Kit (Stemcell technology #19852). T cells were resuspended in lymphocyte media at 6.67×10^5^ cells/ml with an equal number of CD3/28 Dynabeads^TM^ (ThermoFisher #11452D). 10 ng/ml IL-12 and 1 µg/ml IL-4 blocking antibodies were added to T helper 1 cell (T_H_1s) culture and 20 ng/ml IL-4 and 1 µg/ml IFN-γ blocking antibodies were added to T helper 2 cell (T_H_2s) culture. T cells were left at 37°C with 5% CO_2_ for 4 days before use in trans-well assays.

T_H_1s and T_H_2s were subsequently resuspended at 1 x 10^7^/ml in migration media (RPMI1640 with 0.5% fatty-acid free bovine serum albumin, penicillin/streptomyces, and 10mM HEPES buffer). 50 µl T_H_1s and 50 µl T_H_2s were carefully added to the top of each transwell insert at the same time. 10 µl of each were added to a well with 600 µl of media but no transwell insert as the input. The plate was then incubated at 37 °C and 5 % CO_2_ for 3hrs with minimal disturbance.

Afterwards, the transwell inserts were discarded and the cells were transferred to a 96-well round-bottom plate and resuspended in 200 µl of flow buffer. Cells from each well were acquired by flow cytometry for the same amount of time. The input % was calculated by dividing the cell numbers from the migration wells into 5 times cell number from the input well.

### Measurement for GPR183-dependent migration activity in lung tissues

Lung samples were weighed and homogenized in 20 µl times the weight in mg with migration media. The lung homogenate was centrifuged at 300 rpm for 10 min to remove the cellular components first. The supernatant was then transferred to a new tube and centrifuged at 3000 rpm for 10 min to remove other debris. The supernatant was transferred to a new tube and stored at-80°C. M12 cells with 50 % GPR183-GFP transfected (*3*) (gifted by Dr. Jason Cyster) were cultured in Roswell Park Memorial Institute Medium, RPMI, supplemented with 10 % FBS, 10 mM HEPES buffer, penicillin/streptomyces, 2 mM L-Glutamine, and 55 µM 2-Mercaptoethanol at 37 °C and 5 % CO_2_.

M12 cells were resuspended at 1 x 10^7^/ml in migration media. Lung extracts were diluted 1:10 in migration media. 600 µl of media, 10 nM 7 α, 25-HC, or diluted lung extracts were added to wells of a 24-well plate in duplicate and 5 µm transwell inserts were placed on top. 100µl of EBI2-GFP M12 cells (1 x 10^6^) were carefully added to the top of each transwell insert. 20µl of EBI2-GFP M12 cells were added to a well with 600 µl of media but no transwell insert for the input. The plate was then incubated at 37 °C and 5 % CO_2_ for 3 hours with minimal disturbance. Afterwards, the transwell inserts were discarded and the cells were transferred to a 96-well round-bottom plate and resuspended in 200µl of flow buffer for the same analysis by flow cytometry as mentioned above.

### RNA extraction and real-time quantitative PCR (RT-qPCR)

The whole process was performed on ice or 4 °C. Lung samples were homogenized in TRIzol (Invitrogen #15596026) followed by adding 200 µl Chloroform. Samples were well-mixed and centrifuged at 12,900 g for 15 min. The equal volume of 2-propanol was mixed into the supernatant in a new tube and the samples were precipitated on ice for 10 min followed by 15 min centrifuge at 12,900 g. The supernatant was discarded. The precipitated pellets were washed by 75 % ethanol for twice and air-dried for 5 min. RNA samples were resuspended in nuclease-free water and stored at-80 °C or used for the reverse reaction with SuperScript™ IV Reverse Transcriptase (Thermo Fisher #18090010). Complementary DNA (cDNA) was stored at-20 °C or subsequently used for RT-qPCR with PowerTrack™ SYBR Green Master Mix (Thermo Fisher #A46110). *Ch25h* primers, forward: TGCATCACCAGAACTCGTCC, reverse: GGGAAGTCATAGCCCGAGTG. *Cyp7b1* primer, forward: GGAGCCACGACCCTAGATG, reverse: TGCCAAGATAAGGAAGCCAAC. *Cyp46a1* primer: *Cyp27a1* primer, forward: ATCTGGGTTGGGAAGGTG, reverse: CATTGCTCTCCTTGTGCGATG. The level of mRNA expression was all relative to *Rplp0* (also known as 36B4) expression. *Rplp0* primers, forward: GGGCATCACCACGAAAATCTC, reverse: CTGCCGTTGTCAAACACCT.

### Bone marrow-derived macrophage (BMDM) generation and culture

Bone marrow was flushed out from mouse femurs and tibias and filtered through 70 μm into DMEM medium (10% FBS, 10 mM HEPES buffer, penicillin/streptomyces, 2 mM L-Glutamine). M-CSF was produced by M-CSF-NIH3T3 cell line and added into DMDM medium (10%) to support macrophage differentiation. BMDMs were harvested 7 days after differentiation.

### Thioglycolate-elicited peritoneal macrophage (PEM) collection

Thioglycolate (3%) was injected intraperitoneally 4 days before the collection. 10 ml PBS was injected into peritoneal cavity to collect PEMs. PEMs were maintained in DMEM medium (10% FBS, 10 mM HEPES buffer, penicillin/streptomyces, 2 mM L-Glutamine).

### Single-cell RNA sequencing (scRNA-seq) for *Cryptococcus neoformans (Cn)* infection

Mice at different time points were infected separately and harvested at the same time. Single cell suspension was made in the same way as flow cytometry experiments. Hashtag antibodies (TotalSeq™-C0301 to C315 anti-mouse Hashtag 1 to 15 Antibody, Biolegend #155861 to #155889) were added into the staining cocktail. Macrophages (Live/CD45^+^/IV^-^/B220^-^/CD90.2^-^/Ly6G^-^/Ly6C^-^/MertK^+^/CD64^+^) were sorted from these samples and pooled together for generating library. The library was prepared based on Chromium Next GEM Single Cell 5’ Reagent Kits v2 (10X Genomics, document number: CG000220 Rev F). The sequencing was done at Center for Human Immunology/NIAID with NextSeq 1000/2000 P2 XLEAP-SBS Reagent Kit (Illumina #20100987). The parameters were Read 1/Index 1/Index 2/Read 2: 27 / 10 / 10 / 91 cycles. The 10X CellRanger (v7.1.0) mkfastq and count pipelines were respectively used to generate FASTQ files and count matrices with the mm10 reference (refdata-gex-mm10-2020-A). Downstream analysis was performed in R (v4.3.0) using Seurat (v4.3.0) (*4*). Libraries were filtered by applying thresholds based on log-transformed values. Specifically, cells with low total RNA content or a low number of detected genes—both below the 3 median absolute deviations (MAD) lower threshold—were excluded. Additionally, cells with high mitochondrial gene expression levels, high total RNA, high number of cells or high HTO counts, exceeding the 3 MAD upper threshold, were also excluded. After merging the samples using Seurat’s inbuilt merge function, hashtags were demultiplexed using HTODemux from Seurat. Doublets and negatives were removed from further analysis, along with the hashtag HTO-10 (Day10-rep1), which was identified as an outlier based on low expression levels after demultiplexing.

Dimensional reduction of the merged log10 normalized data was performed using 30 principal components and visualized using Uniform Manifold Approximation and Projections (UMAP). Unsupervised clustering was performed with a resolution of 0.7. Cell annotation was performed using SingleR (v2.2.0) (*5*) and mouse expression data (MouseRNAseqData) (*6*) from Celldex (v1.10.1). Cells labeled macrophages from the main cell types were subset for additional processing. Visualization was performed using Seurat’s visualization tools and ggplot2.

### GPR183 bone marrow overexpression

On day 1, donor mice were injected with 150 mg/kg 5-fluorouracil (5-FU) intraperitoneally. On day 4, PLAT-E cells were transfected with MSCV-IRES-*Gfp* or MSCV-*Gpr183*-IRES-*Gfp* vectors to produce mouse-tropic retrovirus. On day 5, bone marrow was collected from the 5-FU-treated donor mice and plated into 24 well-plates with bone marrow culture medium (DMEM with 15 % FBS, 20 ng/mL IL-3, 50 ng/mL IL-6, 100 ng/mL SCF, Pen/Strep, HEPES). On day 6, supernatant was collected from transfected PLAT-E cells and filtered through a 0.45 µm syringe filter. Filtered virus was complemented with 4 µg/ml polybrene and HEPES. Bone marrow cells were then treated with virus and centrifuged at room temperature for 2 hours at 2450 rpm. Bone marrow cells were recovered for 24 hours followed by the second spin-infection and another 24 hours recovery. On day 8, bone marrow cells were collected and washed for transfer into irradiated recipients.

### Thick-tissue imaging

Mice were euthanized and perfused with 20 ml PBS followed by 20 ml 4 % PFA. Inflation with 2 % low-melting agarose was then performed for thick tissue sectioning using a Vibrating Blade Microtome. 500 µm sections were made and processed with Ce3D Tissue Clearing buffer set (*7*) (BioLegend #427702) according to commercial instructions. Briefly, sections were permeabilized at room temperature for 2 days with gentle shaking followed by antibody staining for another 2 days with gentle shaking. Sections were washed 3 times by washing buffer during next 24 hours and then cleared by clearing solution for overnight. Cleared sections were mounted with clearing solution in 4 spacers (120 µm thickness for each) on slides for imaging by Leica SP8 conjugated with 690 nm laser at Bioimaging Research Technologies Branch at NIAID.

### scRNA-seq for *Cn gcs1Δ* infection

Myeloid cells (Live/CD45^+^/IV^-^/B220^-^/CD90.2^-^/Ly-6G^-^/MHCII^+^/CD11c^+^/MHCII^+^/CD64^+^) were sorted at 35 days post-infection. Cells were also labeled by Hashtag antibodies and then pooled for library preparation and sequencing followed the same protocol as mentioned above. Data analysis was performed similarly, and no sample was excluded due to quality issue.

### RNA scope

ACD RNAscope Multiplex Fluorescent V2 assays was used for detection of *Ch25h* and *Cyp7b1*

RNA expression and co-detection of CD11c protein expression. Briefly, mice were euthanized and perfused with 20 ml PBS followed by 20 ml 4 % PFA. OCT was used to inflate lungs through the trachea, and the tissues were flash frozen in molds and sectioned into 8 µm slices. Sections were fixed with 4% PFA and dehydrated for hydrogen peroxide incubation. Sections were then rehydrated for CD11c staining overnight and subsequently crosslinked with 10% neutral buffered formalin. Hybridization of both probes in different channels following the protease treatment was performed, and signal development for two channels was done separately with two different fluorophores. DAPI staining was performed 30 sec before mounting.

---

**Fig. S1.**
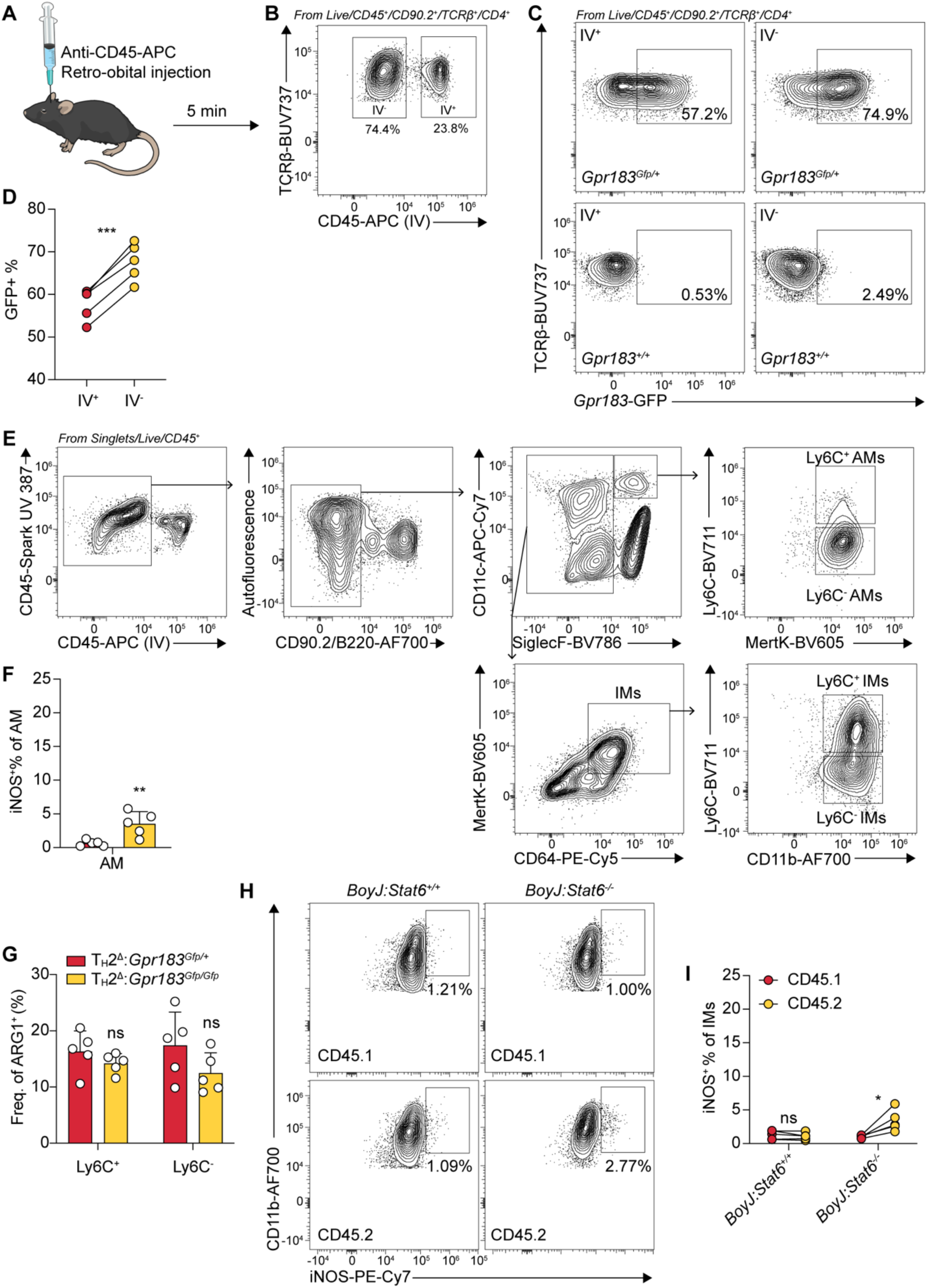
T_H_2 sensing of oxysterols suppress iNOS expression in Ly6C+ macrophages. (A) Schematic of intravenous (IV) CD45 labeling. (B) Representative flow cytometry plot showing IV labeling within the CD4^+^ T cell compartment from *Cn*-infected mouse lung tissues at 10 dpi. (C) Representative flow cytometry plots for *Gpr183*-GFP expression in IV^+^ and IV^-^ CD4^+^ T cells from *Cn*-infected mouse lung tissues at 10 dpi. (D) Quantification of *Gpr183*-GFP^+^ portion in IV^+^ versus IV^-^ CD4^+^ T cells from *Cn*-infected mouse lung tissues at 10 dpi; lines connect paired populations from the same host. \*\*\**P < 0.001*, paired two-tailed Student’s t-test. (E) Gating strategy for lung resident Ly6C^+^ or Ly6C^-^ alveolar macrophages (AMs) and Ly6C^-^ interstitial macrophages (IMs). (F) Frequency of iNOS^+^ cells in lung AMs from *Cn*-infected *Gpr183^TH211^* mice and their controls at 10 dpi. (G) Frequency of ARG1^+^ cells in lung CD64^+^ macrophages from *Cn*-infected *Gpr183^TH211^* mice and their controls at 10 dpi. (H) Representative flow cytometry plots of iNOS expression in CD64^+^ lung macrophages from CD45.1^+^ or CD45.2^+^ donors in STAT6-knockout bone marrow mixed chimera at 10 dpi. (I) Frequency of iNOS+ cells in lung CD64+ macrophages from CD45.1^+^ or CD45.2^+^ donors in STAT6-knockout bone marrow mixed chimera at 10 dpi.

**Fig. S2.**
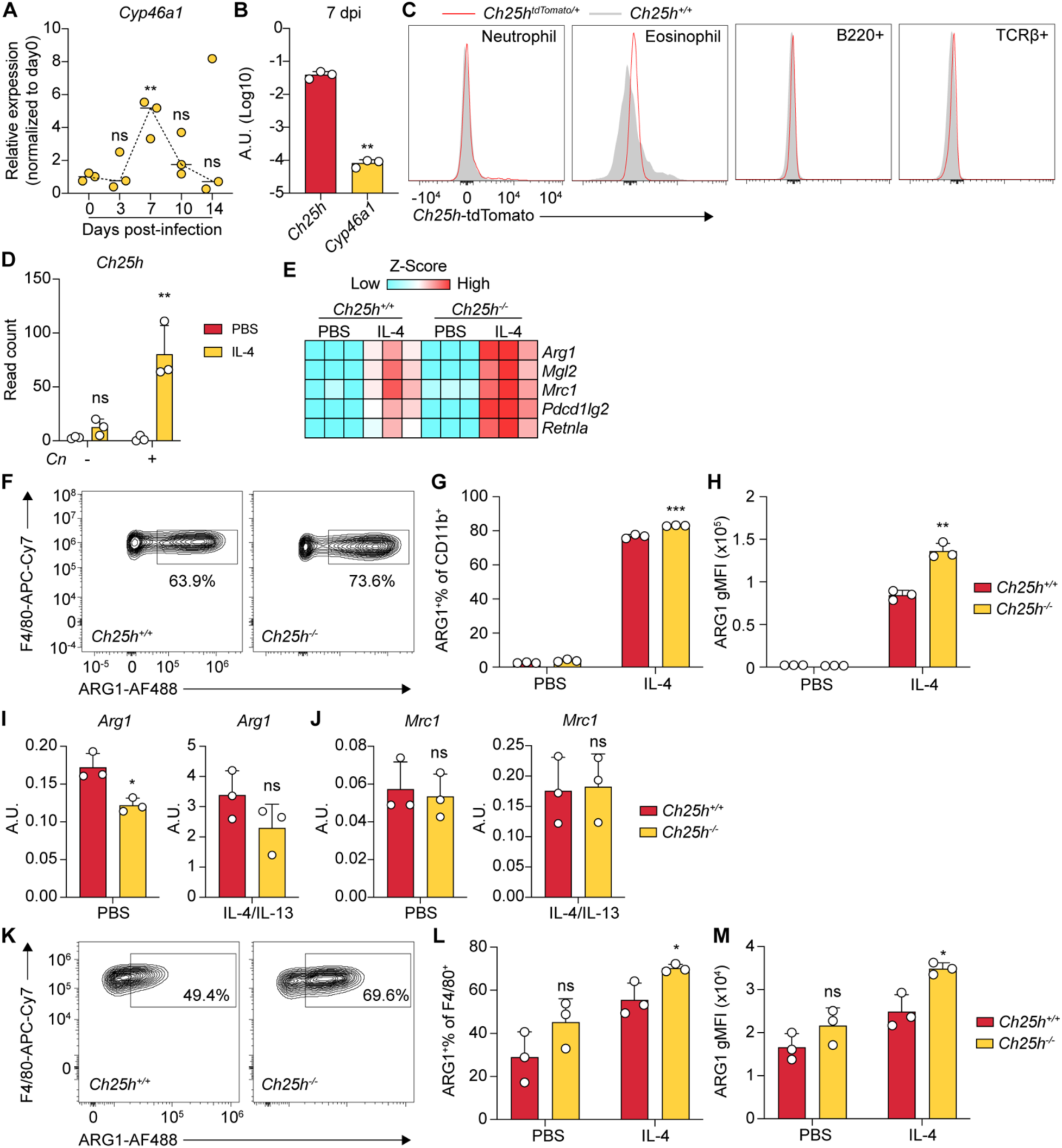
Lung macrophages establish GPR183 chemotactic gradients via *Ch25h* induction. (A) Relative mRNA levels of *Cyp46a1* in lung tissues over time following infection. Expression values were normalized to day 0 to show fold induction. \*\**P < 0.01* and ns, not significant, compared to day 0 by one-way ANOVA. (B) Relative mRNA levels of *Ch25h* and *Cyp46a1* to *Rplp0* in lung tissues from *Cn*-infected mice at 7 dpi. A.U., arbitrary units. \*\**P < 0.01*, unpaired two-tailed Student’s t-test. (C) Representative histograms of *Ch25h*-tdTomato expression across indicated cell populations. All these populations were gated from Live/CD45^+^/IV^-^. Neutrophils were gated by Ly-6G^+^CD11b^+^. Eosinophils were gated by SiglecF^+^CD11b^+^SSC-A^hi^. (D) Read count of *Ch25h* mRNA by bulk RNA-sequencing on bone marrow-derived macrophages (BMDMs) treated by indicated conditions. IL-4, 20 ng/ml; *Cn*, MOI=1. \*\**P < 0.01*; ns, not significant; unpaired two-tailed Student’s t-test. (E) Heatmap of mRNA expression level of indicated markers for alternative activation of macrophages by bulk RNA-sequencing in BMDMs. Expression level was presented as Z-score. IL-4, 20 ng/ml. (F) Representative flow cytometry plot showing ARG1 expression in *Ch25h^+/+^* versus *Ch25h^-/-^* BMDMs after 24 hours-IL-4 treatment. IL-4, 20 ng/ml. (G) Frequency of ARG1^+^ in *Ch25h^+/+^* versus *Ch25h^-/-^* BMDMs after 24 hours-IL-4 or PBS treatment. \*\*\**P < 0.001*, unpaired two-tailed Student’s t-test. (H) Geometric mean fluorescence intensity (gMFI) of ARG1 in *Ch25h^+/+^* versus *Ch25h^-/-^* BMDMs after 24 hours-IL-4 or PBS treatment. \*\**P < 0.01*, unpaired two-tailed Student’s t-test. (I) Relative mRNA levels of *Arg1* to *Rplp0* in thioglycolate-elicited peritoneal macrophages (PEMs) from *Ch25h^+/+^* versus *Ch25h^-/-^* mice after 24 hours-IL-4/IL-13 or PBS treatment. IL-4, 20 ng/ml; IL-13, 20 ng/ml. \**P < 0.05*, ns, not significant, unpaired two-tailed Student’s t-test. (J) Relative mRNA levels of *Mrc1* to *Rplp0* in PEMs from *Ch25h^+/+^* versus *Ch25h^-/-^* mice after 24 hours-IL-4/IL-13 or PBS treatment. ns, not significant; unpaired two-tailed Student’s t-test. (K) Representative flow cytometry plot showing ARG1 expression in *Ch25h^+/+^* versus *Ch25h^-/-^* PEMs after 24 hours-IL-4 treatment. IL-4, 20 ng/ml. (L) Quantification of ARG1^+^ frequency in *Ch25h^+/+^* versus *Ch25h^-/-^* PEMs after 24 hours-IL-4 or PBS treatment. \**P < 0.05*, ns, not significant, unpaired two-tailed Student’s t-test. (M) Quantification of ARG1 gMFI in *Ch25h^+/+^* versus *Ch25h^-/-^* PEMs after 24 hours-IL-4 or PBS treatment. \**P < 0.05*, ns, not significant, unpaired two-tailed Student’s t-test.

**Fig. S3.**
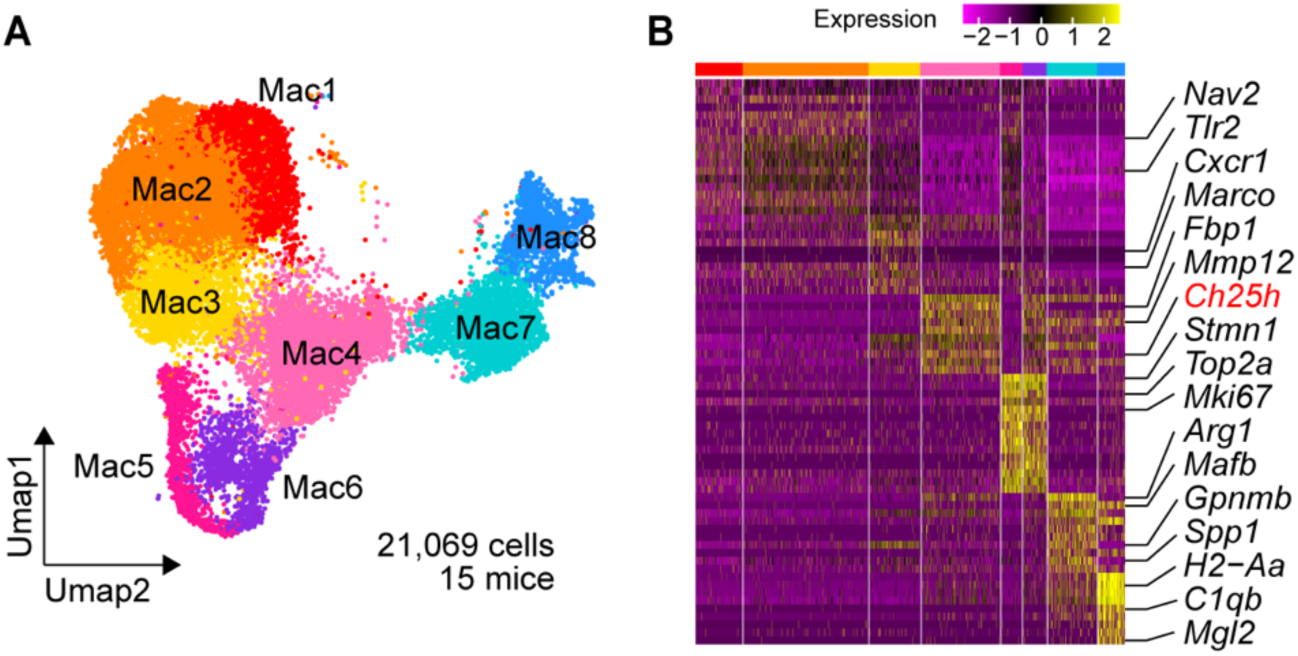
Single cell RNA-sequencing on Ly6C^-^ lung macrophages. (A) UMAP visualization of original macrophage clusters from combined scRNA-seq replicates. (B) Heatmap of top 10 differential expressed genes in each cluster.

**Fig. S4.**
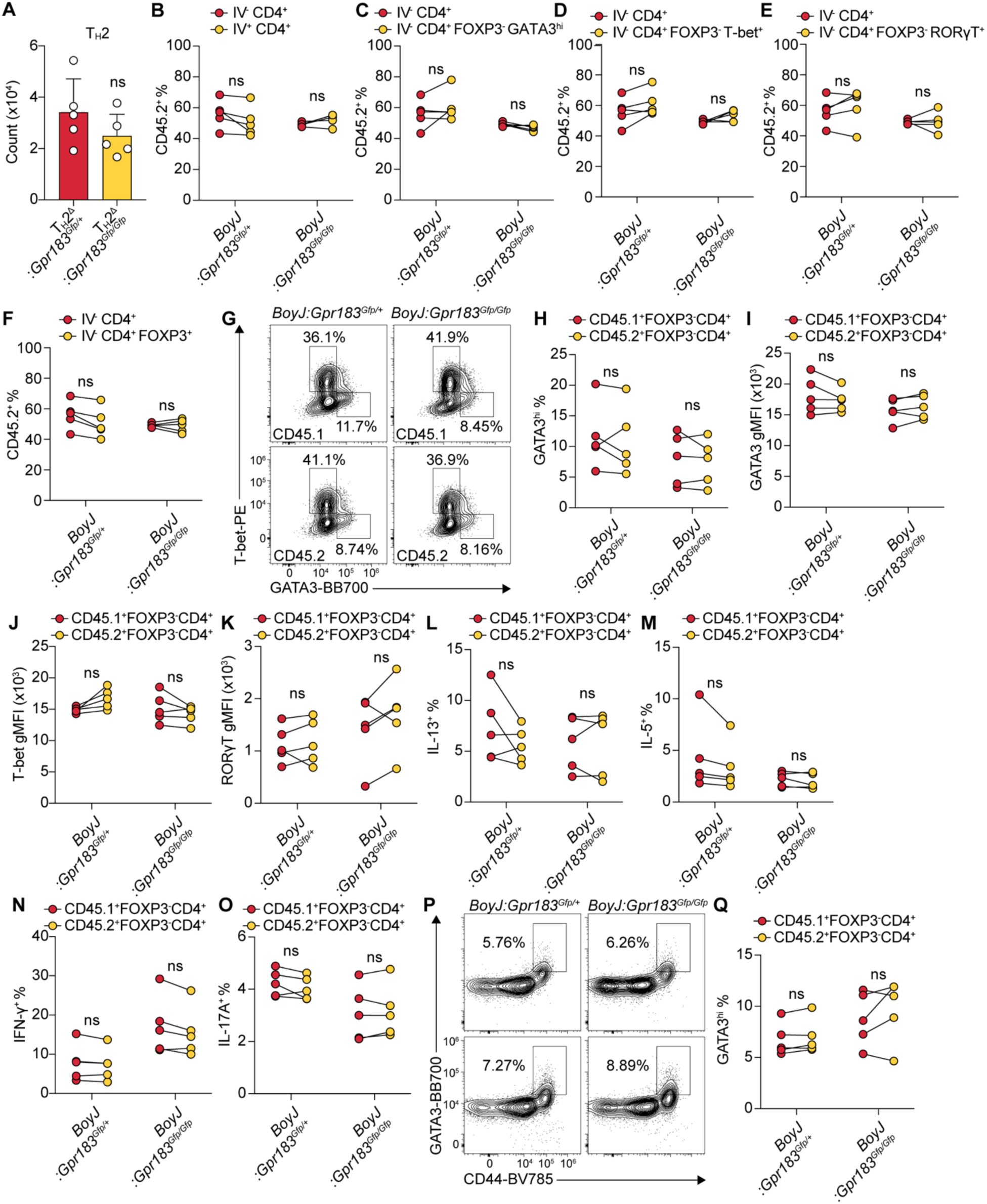
GPR183 is dispensable for T cell recruitment or T_H_2 differentiation. (A) Quantification of lung T_H_2 cells in *Cn*-infected *Gpr183^TH211^* mice and their controls at 10 dpi. Lung T_H_2 cells were gated by Live/CD45^+^/IV^-^/CD90.2^+^/TCRb^+^/CD4^+^/FOXP3^-^/T-bet^-^/GATA3^hi^. (B) Proportional contribution of CD45.2^+^ cells to IV^+^ versus IV^-^CD4^+^ T cells in lungs from *Cn*-infected mice with GPR183-knockout bone marrow mixed chimera and their controls at 10 dpi. ns, not significant by paired two-tailed Student’s t-test. (C) Proportional contribution of CD45.2^+^ cells to total IV^-^ CD4^+^ T cells versus IV^-^ CD4^+^ FOXP3^-^ GATA3^hi^ T_H_2 subset in lungs from *Cn*-infected mice with GPR183-knockout bone marrow mixed chimera and their controls at 10 dpi. ns, not significant by paired two-tailed Student’s t-test. (D) Proportional contribution of CD45.2^+^ cells to IV^-^ CD4^+^ T cells and IV^-^ CD4^+^ FOXP3^-^T-bet^+^ T_H_1 cells from *Cn*-infected mice with GPR183-knockout bone marrow mixed chimera and their controls at 10 dpi. ns, not significant by paired two-tailed Student’s t-test. (E) Proportional contribution of CD45.2^+^ cells to IV^-^ CD4^+^ T cells and IV^-^ CD4^+^ FOXP3^-^ RORγT^+^ T_H_17 cells from *Cn*-infected mice with GPR183-knockout bone marrow mixed chimera and their controls at 10 dpi. ns, not significant by paired two-tailed Student’s t-test. (F) Proportional contribution of CD45.2^+^ cells to IV^-^ CD4^+^ T cells and IV^-^ CD4^+^ FOXP3^+^ T_reg_ cells from *Cn*-infected mice with GPR183-knockout bone marrow mixed chimera and their controls at 10 dpi. ns, not significant by paired two-tailed Student’s t-test. (G) Representative flow cytometry plots of T-bet and GATA3 expression in CD45.1^+^ and CD45.2^+^ FOXP3^-^CD4^+^ T cell population in the lungs from *Cn*-infected mice with GPR183-knockout bone marrow mixed chimera and their controls at 10 dpi. Gating: Singlets/Live/CD45^+^/IV^-^/CD90.2^+^/TCR^+^/CD4^+^/FOXP3^-^. (H) Frequency of GATA3^hi^ cells within CD45.1^+^ or CD45.2^+^ FOXP3^-^ CD4^+^ T cells in lungs from *Cn*-infected mice with GPR183-knockout bone marrow mixed chimera and their controls at 10 dpi. ns, not significant by paired two-tailed Student’s t-test. (I) GATA3 gMFI in CD45.1^+^ or CD45.2^+^ FOXP3^-^ CD4^+^ T cells from lungs in *Cn*-infected mice with GPR183-knockout bone marrow mixed chimera and their controls at 10 dpi. ns, not significant by paired two-tailed Student’s t-test. (J) T-bet geometric mean of fluorescent intensity (gMFI) in CD45.1^+^ or CD45.2^+^ FOXP3^-^ CD4^+^ T cells from lungs in *Cn*-infected mice with GPR183-knockout bone marrow mixed chimera and their controls at 10 dpi. ns, not significant by paired two-tailed Student’s t-test. (K) RORγT gMFI in CD45.1^+^ or CD45.2^+^ FOXP3^-^ CD4^+^ T cells from lungs in *Cn*-infected mice with GPR183-knockout bone marrow mixed chimera and their controls at 10 dpi. ns, not significant by paired two-tailed Student’s t-test. (L) Frequency of IL-13^+^ cells among CD45.1^+^ and CD45.2^+^ FOXP3^-^ CD4^+^ T cells in lungs from *Cn*-infected mice with GPR183-knockout bone marrow mixed chimera and their controls at 10 dpi. ns, not significant by paired two-tailed Student’s t-test. (M) Frequency of IL-5^+^ cells among CD45.1^+^ and CD45.2^+^ FOXP3^-^ CD4^+^ T cells in lungs from *Cn*-infected mice with GPR183-knockout bone marrow mixed chimera and their controls at 10 dpi. ns, not significant by paired two-tailed Student’s t-test. (N) Frequency of IFNγ^+^ cells among CD45.1^+^ and CD45.2^+^ FOXP3^-^ CD4^+^ T cells in lungs from *Cn*-infected mice with GPR183-knockout bone marrow mixed chimera and their controls at 10 dpi. ns, not significant by paired two-tailed Student’s t-test. (O) Frequency of IL-17^+^ cells among CD45.1^+^ and CD45.2^+^ FOXP3^-^ CD4^+^ T cells in lungs from *Cn*-infected mice with GPR183-knockout bone marrow mixed chimera and their controls at 10 dpi. ns, not significant by paired two-tailed Student’s t-test. (P) Representative flow cytometry plots of GATA3 expression in mediastinal lymph node CD45.1^+^ and CD45.2^+^ FOXP3^-^CD4^+^ T cells from *Cn*-infected mice with GPR183-knockout bone marrow mixed chimera and their controls at 10 dpi. Gating: Singlets/Live/CD45^+^/IV^-^/CD90.2^+^/TCR^+^/CD4^+^/FOXP3^-^. (Q) Frequency of GATA3^hi^ cells within CD45.1^+^ and CD45.2^+^ FOXP3^-^ CD4^+^ T cells in mediastinal lymph nodes from *Cn*-infected mice with GPR183-knockout bone marrow mixed chimera and their controls at 10 dpi. ns, not significant by paired two-tailed Student’s t-test.

**Fig. S5.**
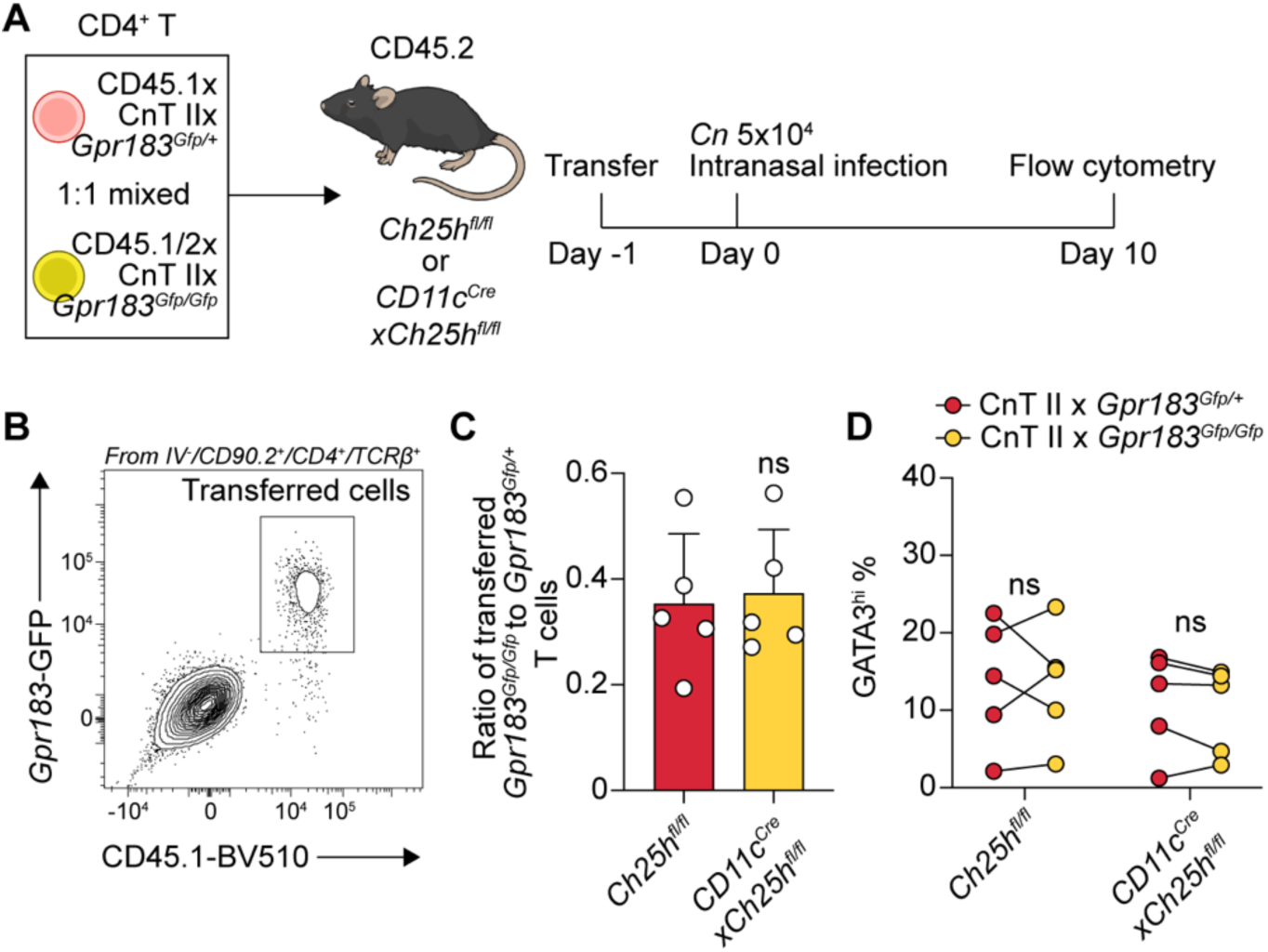
GPR183 is dispensable for cryptococcal antigen-specific T cell recruitment or T_H_2 differentiation. (A) Schematic for adoptive T cell transfer experiment. Cryptococcal-specific TCR transgenic (CnT II) *Gpr183^Gfp/+^*or *GPR183^Gfp/Gfp^* CD4^+^ T cells were transferred into *Ch25h^fl/fl^* or *CD11c^Cre^xCh25h^fl/fl^* mice one day before infection (-1 dpi). Flow analysis was performed at 10 dpi. (B) Representative flow cytometry plot for transferred *Gpr183*-GFP^+^ CnT II T cells. (C) Ratio of transferred CnT II *Gpr183^Gfp/+^* to *GPR183^Gfp/Gfp^* T cells in lungs of *Ch25h^fl/fl^* and *CD11c^Cre^*x*Ch25h^fl/fl^* mice at 10 dpi. ns, not significant by unpaired two-tailed Student’s t-test. (D) Frequency of GATA3^hi^ cells within transferred CnT II *Gpr183^Gfp/+^* or *GPR183^Gfp/Gfp^* FOXP3^-^ CD4^+^ T cells in lung tissues from *Cn*-infected *Ch25h^fl/fl^* or *CD11c^Cre^xCh25h^fl/fl^* mice at 10 dpi. ns, not significant by paired two-tailed Student’s t-test.

**Fig. S6.**
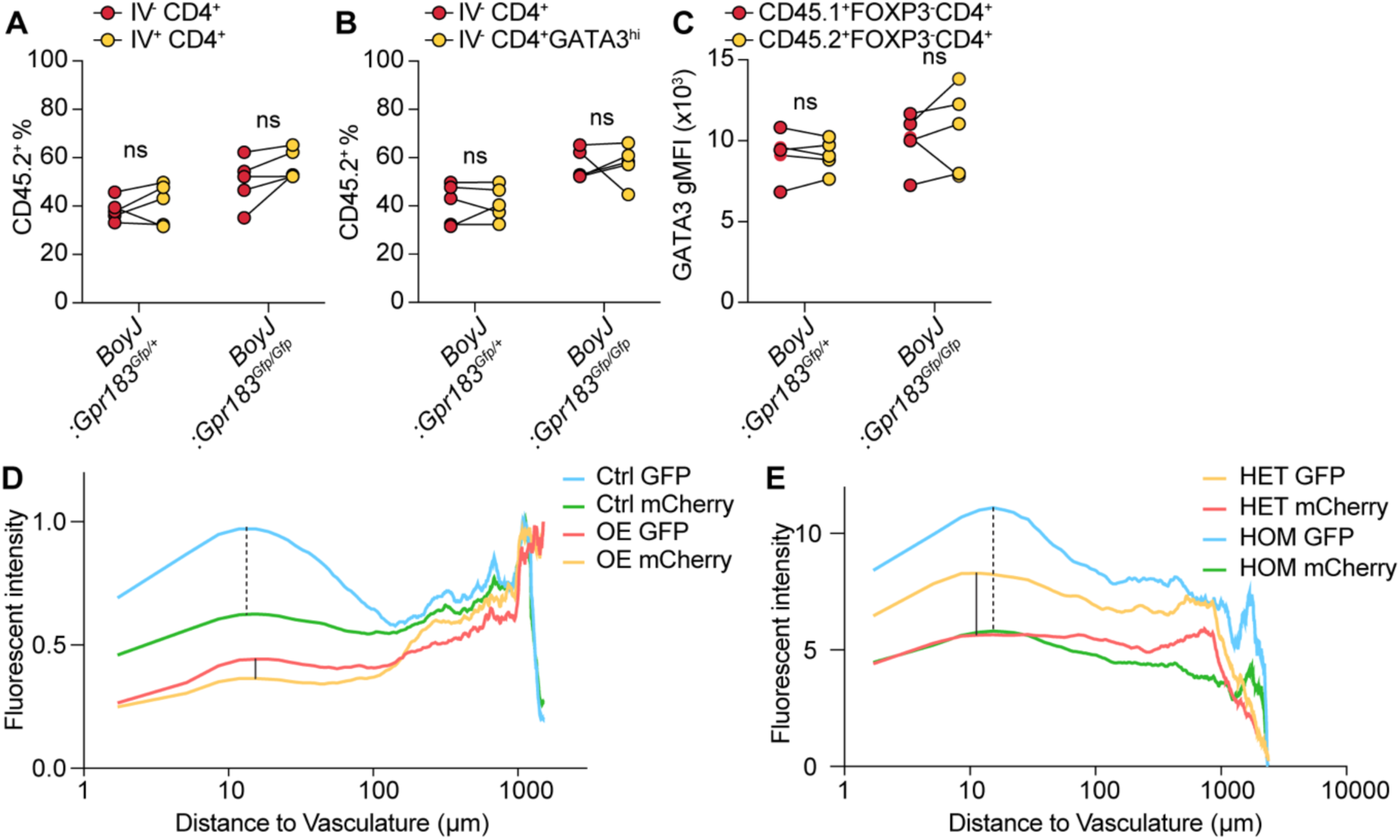
GPR183 guides TH2 intra-pulmonary positioning. (A) Proportional contribution of CD45.2^+^ cells to IV^-^ versus IV^+^ CD4^+^ T cells from *Cn gcs1Δ*-infected mice with GPR183-knockout bone marrow mixed chimera and their controls at 21 dpi. (B) Proportional contribution of CD45.2^+^ cells to total IV^-^ CD4^+^ T cells versus IV^-^ CD4^+^ FOXP3^-^ GATA3^hi^ T_H_2 cells from *Cn gcs1Δ*-infected mice with GPR183-knockout bone marrow mixed chimera and their controls 21 dpi. (C) GATA3 gMFI in CD45.1^+^ or CD45.2^+^ FOXP3^-^ CD4^+^ T cells from lungs in *Cn gcs1Δ*-infected mice with GPR183-knockout bone marrow mixed chimera and their controls 21 dpi. (D) Quantification of fluorescent intensity at indicated distance to vasculature from images in Fig. 4A. (E) Quantification of fluorescent intensity at indicated distance to vasculature from images in Fig. 4B.

**Fig. S7.**
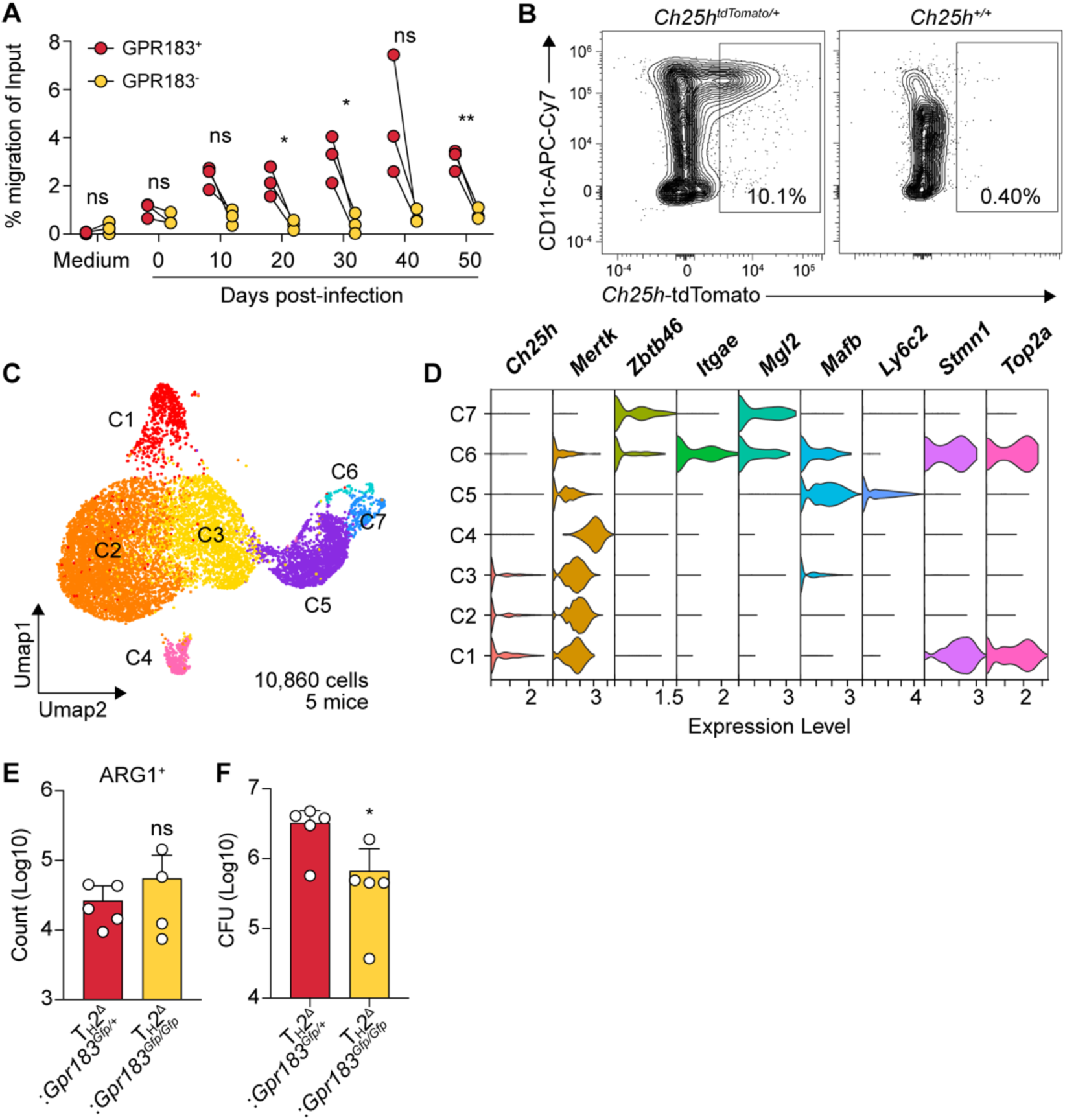
Granuloma-associated macrophages antagonize fungal clearance by positioning T_H_2 cells. (A) Quantification of migration activity of GPR183^+^ versus GPR183^-^ M12 cells in response to lung extracts collected at indicated time points or migration media alone. \**P < 0.05*, \*\**P < 0.01*; ns, not significant; paired two-tailed Student’s t-test. (B) Representative flow cytometry plots of *Ch25h*-tdTomato expression in CD45^+^ IV^-^ immune cells at 35 dpi. (C) UMAP visualization of original myeloid clusters from combined scRNA-seq replicates. (D) Violin plots of the selected lineage marker expression across myeloid clusters. (E) Frequency of ARG1^+^ cells in lung CD64^+^ macrophages from *Cn gcs1Δ*-infected *Gpr183^TH211^* mice and their controls at 21 dpi. (F) Pulmonary fungal burden in *Cn gcs1Δ*-infected *Gpr183^TH211^* mice and their controls at 35 dpi. CFU, colony forming unit.

**Table S1.**

| **Antibodies** |  |  |
| --- | --- | --- |
| AF488 anti-mouse ARG1 (clone A1EXF5) | eBioscience | Cat# 53-3697-80 |
| AF488 anti-GFP (polyclonal) | ThermoFisher | Cat# A-21311 |
| AF647 anti-mouse Ly-6G (clone 1A8) | BioLegend | Cat# 127609 |
| AF700 anti-mouse EpCAM (clone G8.8) | BioLegend | Cat# 118240 |
| APC anti-mouse CD45 (clone 30-F11) | BioLegend | Cat# 103112 |
| APC-Cy7 anti-mouse CD11c (clone N418) | BioLegend | Cat# 117324 |
| APC-Cy anti-mouse IL-17A (clone TC11-18H10.1) | BioLegend | Cat# 506940 |
| BB700 anti-mouse GATA3 (clone L50823) | BD<br>Bioscience | Cat# 566643 |
| BUV737 anti-mouse TCRβ (clone H57-597) | BD<br>Bioscience | Cat# 612821 |
| BV421 anti-mouse CD3 (clone 17A2) | BioLegend | Cat# 100227 |
| BV421 anti-mouse IL-5 (clone TRFK5) | BioLegend | Cat# 504311 |
| BV421 anti-mouse Foxp3 (clone MF-14) | BioLegend | Cat# 126419 |
| BV421 anti-mouse PD-L2 (clone TY25) | BioLegend | Cat# 107219 |
| BV510 anti-mouse CD103 (clone 2E7) | BioLegend | Cat# 121423 |
| BV510 anti-mouse CD45.1 (clone A20) | BD<br>Bioscience | Cat# 565278 |
| BV510 anti-mouse CD90.2 (clone 30-H12) | BioLegend | Cat# 105335 |
| BV605 anti-mouse CD4 (clone GK1.5) | BioLegend | Cat# 100451 |
| BV605 anti-mouse MertK (clone 2B10C42) | BioLegend | Cat# 151517 |
| BV650 anti-mouse Ki67 (clone 11F6) | BioLegend | Cat# 151215 |
| BV711 anti-mouse Ly6C (clone HK1.4) | BioLegend | Cat# 128037 |
| BV711 anti-mouse IFNγ (clone XMG1.2) | BD<br>Bioscience | Cat# 563736 |
| BV786 anti-mouse CD44 (clone IM7) | BD<br>Bioscience | Cat# 563736 |
| BV786 anti-mouse SiglecF (clone E50-2440) | BD<br>Bioscience | Cat# 740956 |
| FITC anti-mouse CD8 (clone S18018E) | BioLegend | Cat# 162313 |
| PE anti-mouse T-bet (clone 4B10) | BioLegend | Cat# 644809 |
| PE-Cy5 anti-mouse CD64 (clone X54-5/7.1) | BioLegend | Cat# 139332 |
| PE-Cy7 anti-mouse CD11c (clone N418) | BioLegend | Cat# 117317 |
| PE-Cy7 anti-mouse iNOS (clone CXNFT) | eBioscience | Cat# 25-5920-80 |
| PE-Cy7 anti-mouse Ly6G (clone 1A8) | BioLegend | Cat# 127617 |
| PE-Cy7 anti-mouse CD301b (MGL2) (clone URA-1) | BioLegend | Cat# 146807 |
| PE-Cy7 anti-mouse RORγt (clone B2D) | eBioscience | Cat# 25-6981-82 |
| PerCP-Cy5.5 anti-mouse MHCII (clone M5/114.15.2) | BioLegend | Cat# 107625 |
| R718 anti-mouse IL-13 (clone W19-895) | BD<br>Bioscience | Cat# 569945 |
| Spark UV 387 anti-mouse CD45 (clone 30F-11) | BioLegend | Cat# 103188 |
| Spark UV 387 anti-mouse CD45.2 (clone 104) | BioLegend | Cat# 109868 |

## References

1. E. Lu, E. V. Dang, J. G. McDonald, J. G. Cyster, Distinct oxysterol requirements for positioning naïve and activated dendritic cells in the spleen. Sci Immunol 2, (2017).

2. J. Li, E. Lu, T. Yi, J. G. Cyster, EBI2 augments Tfh cell fate by promoting interaction with IL-2-quenching dendritic cells. Nature 533, 110–114 (2016).

3. T. Yi, J. G. Cyster, EBI2-mediated bridging channel positioning supports splenic dendritic cell homeostasis and particulate antigen capture. Elife 2, e00757 (2013).

4. L. M. Kelly, J. P. Pereira, T. Yi, Y. Xu, J. G. Cyster, EBI2 guides serial movements of activated B cells and ligand activity is detectable in lymphoid and nonlymphoid tissues. J Immunol 187, 3026–3032 (2011).

5. J. Emgård et al., Oxysterol Sensing through the Receptor GPR183 Promotes the Lymphoid-Tissue-Inducing Function of Innate Lymphoid Cells and Colonic Inflammation. Immunity 48, 120–132.e128 (2018).

6. C. Chu et al., Anti-microbial Functions of Group 3 Innate Lymphoid Cells in Gut-Associated Lymphoid Tissues Are Regulated by G-Protein-Coupled Receptor 183. Cell Rep 23, 3750–3758 (2018).

7. K. Y. Chen et al., Inflammation switches the chemoattractant requirements for naive lymphocyte entry into lymph nodes. Cell 188, 1019–1035.e1022 (2025).

8. S. Ceglia et al., An epithelial cell-derived metabolite tunes immunoglobulin A secretion by gut-resident plasma cells. Nat Immunol 24, 531–544 (2023).

9. M. Frascoli et al., Skin γδ T cell inflammatory responses are hardwired in the thymus by oxysterol sensing via GPR183 and calibrated by dietary cholesterol. Immunity 56, 562–575.e566 (2023).

10. F. Wanke et al., EBI2 Is Highly Expressed in Multiple Sclerosis Lesions and Promotes Early CNS Migration of Encephalitogenic CD4 T Cells. Cell Rep 18, 1270–1284 (2017).

11. T. Yi et al., Oxysterol gradient generation by lymphoid stromal cells guides activated B cell movement during humoral responses. Immunity 37, 535–548 (2012).

12. S. Hannedouche et al., Oxysterols direct immune cell migration via EBI2. Nature 475, 524–527 (2011).

13. C. Liu et al., Oxysterols direct B-cell migration through EBI2. Nature 475, 519–523 (2011).

14. J. G. Cyster, E. V. Dang, A. Reboldi, T. Yi, 25-Hydroxycholesterols in innate and adaptive immunity. Nature Reviews Immunology 14, 731–743 (2014).

15. E. V. Dang, A. Reboldi, Cholesterol sensing and metabolic adaptation in tissue immunity. Trends Immunol 45, 861–870 (2024).

16. A. P. Baptista et al., The Chemoattractant Receptor Ebi2 Drives Intranodal Naive CD4+ T Cell Peripheralization to Promote Effective Adaptive Immunity. Immunity 50, 1188–1201.e1186 (2019).

17. S. Hannedouche et al., Oxysterols direct immune cell migration via EBI2. Nature 475, 524–527 (2011).

18. J. Li, E. Lu, T. Yi, J. G. Cyster, EBI2 augments Tfh cell fate by promoting interaction with IL-2-quenching dendritic cells. Nature 533, 110–114 (2016).

19. L. B. Rodda et al., Single-Cell RNA Sequencing of Lymph Node Stromal Cells Reveals Niche-Associated Heterogeneity. Immunity 48, 1014–1028.e1016 (2018).

20. U. M. Gundra et al., Alternatively activated macrophages derived from monocytes and tissue macrophages are phenotypically and functionally distinct. Blood 123, e110–e122 (2014).

21. K. Park, A. L. Scott, Cholesterol 25-hydroxylase production by dendritic cells and macrophages is regulated by type I interferons. J Leukoc Biol 88, 1081–1087 (2010).

22. D. R. Bauman et al., 25-Hydroxycholesterol secreted by macrophages in response to Toll-like receptor activation suppresses immunoglobulin A production. Proc Natl Acad Sci U S A 106, 16764–16769 (2009).

23. T. Zou, O. Garifulin, R. Berland, V. L. Boyartchuk, Listeria monocytogenes infection induces prosurvival metabolic signaling in macrophages. Infect Immun 79, 1526–1535 (2011).

24. J. Xiao et al., 25-Hydroxycholesterol regulates lysosome AMP kinase activation and metabolic reprogramming to educate immunosuppressive macrophages. Immunity 57, 1087–1104.e1087 (2024).

25. E. V. Dang, J. G. McDonald, D. W. Russell, J. G. Cyster, Oxysterol Restraint of Cholesterol Synthesis Prevents AIM2 Inflammasome Activation. Cell 171, 1057–1071.e1011 (2017).

26. A. Reboldi et al., 25-Hydroxycholesterol suppresses interleukin-1–driven inflammation downstream of type I interferon. Science 345, 679–684 (2014).

27. E. S. Gold et al., 25-Hydroxycholesterol acts as an amplifier of inflammatory signaling. Proceedings of the National Academy of Sciences 111, 10666–10671 (2014).

28. S. Y. Liu et al., Interferon-inducible cholesterol-25-hydroxylase broadly inhibits viral entry by production of 25-hydroxycholesterol. Immunity 38, 92–105 (2013).

29. M. Blanc et al., The Transcription Factor STAT-1 Couples Macrophage Synthesis of 25-Hydroxycholesterol to the Interferon Antiviral Response. Immunity 38, 106–118 (2013).

30. C. Li et al., 25-Hydroxycholesterol Protects Host against Zika Virus Infection and Its Associated Microcephaly in a Mouse Model. Immunity 46, 446–456 (2017).

31. M. E. Abrams et al., Oxysterols provide innate immunity to bacterial infection by mobilizing cell surface accessible cholesterol. Nature Microbiology 5, 929–942 (2020).

32. Q. D. Zhou et al., Interferon-mediated reprogramming of membrane cholesterol to evade bacterial toxins. Nature Immunology 21, 746–755 (2020).

33. F. O. Martinez et al., Genetic programs expressed in resting and IL-4 alternatively activated mouse and human macrophages: similarities and differences. Blood 121, e57–69 (2013).

34. A. Reboldi et al., Inflammation. 25-Hydroxycholesterol suppresses interleukin-1-driven inflammation downstream of type I interferon. Science 345, 679–684 (2014).

35. M. Blanc et al., The transcription factor STAT-1 couples macrophage synthesis of 25-hydroxycholesterol to the interferon antiviral response. Immunity 38, 106–118 (2013).

36. L. Wirth et al., High-Dimensional Analysis of Type 2 Lymphocyte Dynamics During Mepolizumab or Dupilumab Treatment in Severe Asthma. Allergy n/a.

37. R. L. Gieseck, M. S. Wilson, T. A. Wynn, Type 2 immunity in tissue repair and fibrosis. Nature Reviews Immunology 18, 62–76 (2018).

38. C. M. Lloyd, R. J. Snelgrove, Type 2 immunity: Expanding our view. Science Immunology 3, eaat1604 (2018).

39. B. Spellberg, J. Edwards, Type 1/Type 2 Immunity in Infectious Diseases. Clinical infectious diseases: an official publication of the Infectious Diseases Society of America 32, 76–102 (2001).

40. Y. Zheng, E. V. Dang, Novel mechanistic insights underlying fungal allergic inflammation. PLoS Pathog 19, e1011623 (2023).

41. Y. Zheng et al., Alternatively activated monocyte-derived myeloid cells promote extracellular pathogen persistence within pulmonary fungal granulomas. bioRxiv, 2025.2005.2023.655817 (2025).

42. T. Imai et al., Selective recruitment of CCR4-bearing Th2 cells toward antigen-presenting cells by the CC chemokines thymus and activation-regulated chemokine and macrophage-derived chemokine. International Immunology 11, 81–88 (1999).

43. Z. Mikhak et al., Contribution of CCR4 and CCR8 to antigen-specific T(H)2 cell trafficking in allergic pulmonary inflammation. J Allergy Clin Immunol 123, 67–73.e63 (2009).

44. I. D. Iliev et al., Focus on fungi. Cell 187, 5121–5127 (2024).

45. W. H. Organization, “WHO fungal priority pathogens list to guide research, development and public health action,” (2022).

46. J. P. Pereira, L. M. Kelly, Y. Xu, J. G. Cyster, EBI2 mediates B cell segregation between the outer and centre follicle. Nature 460, 1122–1126 (2009).

47. R. K. Gurram et al., Crosstalk between ILC2s and Th2 cells varies among mouse models. Cell Reports 42, (2023).

48. Z. Czimmerer et al., The Transcription Factor STAT6 Mediates Direct Repression of Inflammatory Enhancers and Limits Activation of Alternatively Polarized Macrophages. Immunity 48, 75–90.e76 (2018).

49. Y. Ohmori, T. A. Hamilton, STAT6 is required for the anti-inflammatory activity of interleukin-4 in mouse peritoneal macrophages. J Biol Chem 273, 29202–29209 (1998).

50. E. V. Dang et al., Secreted fungal virulence effector triggers allergic inflammation via TLR4. Nature 608, 161–167 (2022).

51. G. Bidault et al., SREBP1-induced fatty acid synthesis depletes macrophages antioxidant defences to promote their alternative activation. Nature Metabolism 3, 1150–1162 (2021).

52. M. S. Brown, J. L. Goldstein, The SREBP pathway: regulation of cholesterol metabolism by proteolysis of a membrane-bound transcription factor. Cell 89, 331–340 (1997).

53. Z. Liu et al., Fate Mapping via Ms4a3-Expression History Traces Monocyte-Derived Cells. Cell 178, 1509–1525.e1519 (2019).

54. J. Van den Bossche et al., Alternatively activated macrophages engage in homotypic and heterotypic interactions through IL-4 and polyamine-induced E-cadherin/catenin complexes. Blood 114, 4664–4674 (2009).

55. M. R. Cronan et al., Macrophage Epithelial Reprogramming Underlies Mycobacterial Granuloma Formation and Promotes Infection. Immunity 45, 861–876 (2016).

56. A. C. Bohrer et al., Rapid GPR183-mediated recruitment of eosinophils to the lung after Mycobacterium tuberculosis infection. Cell Rep 40, 111144 (2022).

57. C. X. Foo et al., GPR183 antagonism reduces macrophage infiltration in influenza and SARS-CoV-2 infection. Eur Respir J 61, (2023).

58. T. M. Conlon, A. Yildirim, Oxysterol metabolism dictates macrophage influx during SARS-CoV-2 infection. Eur Respir J 61, (2023).

59. M. D. Ngo et al., A Blunted GPR183/Oxysterol Axis During Dysglycemia Results in Delayed Recruitment of Macrophages to the Lung During Mycobacterium tuberculosis Infection. J Infect Dis 225, 2219–2228 (2022).

60. Z. J. Shen et al., Epstein-Barr Virus-induced Gene 2 Mediates Allergen-induced Leukocyte Migration into Airways. Am J Respir Crit Care Med 195, 1576–1585 (2017).

61. K. Sato et al., Production of IL-17A at Innate Immune Phase Leads to Decreased Th1 Immune Response and Attenuated Host Defense against Infection with Cryptococcus deneoformans. J Immunol 205, 686–698 (2020).

62. M. S. Fu, K. Kawakami, R. A. Drummond, Adoptive Transfer of Cryptococcus neoformans-Specific CD4 T-Cells to Study Anti-fungal Lymphocyte Responses In Vivo. Methods Mol Biol 2667, 99–112 (2023).

63. P. C. Rittershaus et al., Glucosylceramide synthase is an essential regulator of pathogenicity of Cryptococcus neoformans. J Clin Invest 116, 1651–1659 (2006).

64. S. Zhou et al., Pathogenic mycobacterium upregulates cholesterol 25-hydroxylase to promote granuloma development via foam cell formation. iScience 27, 109204 (2024).

65. W. Zeng et al., Characterization of Anti-Interferon-γ Antibodies in HIV-Negative Patients Infected With Disseminated Talaromyces marneffei and Cryptococcosis. Open Forum Infect Dis 6, ofz208 (2019).

66. M. J. Davis et al., Inbred SJL mice recapitulate human resistance to Cryptococcus infection due to differential immune activation. mBio 14, e0212323 (2023).

67. K. Kawakami et al., Contribution of interferon-gamma in protecting mice during pulmonary and disseminated infection with Cryptococcus neoformans. FEMS Immunol Med Microbiol 13, 123–130 (1996).

68. G. H. Chen et al., The gamma interferon receptor is required for the protective pulmonary inflammatory response to Cryptococcus neoformans. Infect Immun 73, 1788–1796 (2005).

69. S. E. Hardison et al., Pulmonary infection with an interferon-gamma-producing Cryptococcus neoformans strain results in classical macrophage activation and protection. Am J Pathol 176, 774–785 (2010).

70. K. Wang et al., Innate cells and STAT1-dependent signals orchestrate vaccine-induced protection against invasive Cryptococcus infection. mBio 15, e0194424 (2024).

71. A. C. Bohrer et al., Rapid GPR183-mediated recruitment of eosinophils to the lung after Mycobacterium tuberculosis infection. Cell Reports 40, 111144 (2022).

72. Y. Lavin et al., Tissue-Resident Macrophage Enhancer Landscapes Are Shaped by the Local Microenvironment. Cell 159, 1312–1326 (2014).

73. L. Zhang et al., GPR183 targets lung-resident CD301b<sup>+</sup> conventional dendritic cells type 2 to a subtissular TSLP – TSLP receptor mediated survival niche within the adventital cuff. bioRxiv, 2022.2008.2028.505379 (2022).

74. E. F. McCaffrey et al., The immunoregulatory landscape of human tuberculosis granulomas. Nature Immunology 23, 318–329 (2022).

75. A. J. Pagán, L. Ramakrishnan, The Formation and Function of Granulomas. Annual Review of Immunology 36, 639–665 (2018).

## References

1. J. P. Pereira, L. M. Kelly, Y. Xu, J. G. Cyster, EBI2 mediates B cell segregation between the outer and centre follicle. Nature 460, 1122–1126 (2009).

2. M. Frascoli et al., Skin γδ T cell inflammatory responses are hardwired in the thymus by oxysterol sensing via GPR183 and calibrated by dietary cholesterol. Immunity 56, 562–575.e566 (2023).

3. J. P. Pereira, L. M. Kelly, Y. Xu, J. G. Cyster, EBI2 mediates B cell segregation between the outer and centre follicle. Nature 460, 1122–1126 (2009).

4. Y. Hao et al., Integrated analysis of multimodal single-cell data. Cell 184, 3573–3587.e3529 (2021).

5. D. Aran et al., Reference-based analysis of lung single-cell sequencing reveals a transitional profibrotic macrophage. Nature Immunology 20, 163–172 (2019).

6. B. A. Benayoun et al., Remodeling of epigenome and transcriptome landscapes with aging in mice reveals widespread induction of inflammatory responses. Genome Res 29, 697–709 (2019).

7. W. Li, R. N. Germain, M. Y. Gerner, High-dimensional cell-level analysis of tissues with Ce3D multiplex volume imaging. Nature Protocols 14, 1708–1733 (2019).

